# Spatial Transcriptional Mapping of the Human Nephrogenic Program

**DOI:** 10.1101/2020.04.27.060749

**Authors:** Nils O. Lindström, Rachel Sealfon, Xi Chen, Riana Parvez, Andrew Ransick, Guilherme De Sena Brandine, Jinjin Guo, Bill Hill, Tracy Tran, Albert D. Kim, Jian Zhou, Alicja Tadych, Aaron Watters, Aaron Wong, Elizabeth Lovero, Brendan H. Grubbs, Matthew E. Thornton, Jill A. McMahon, Andrew D. Smith, Seth W. Ruffins, Chris Armit, Olga G. Troyanskaya, Andrew P. McMahon

## Abstract

Congenital abnormalities of the kidney and urinary tract are amongst the most common birth defects affecting 3% of newborns. The human kidney develops over a 30-week period in which a nephron progenitor pool gives rise to around a million nephrons. To establish a framework for human nephrogenesis, we spatially resolved a stereotypical process by which equipotent nephron progenitors generate a nephron anlagen, then applied data-driven approaches to construct three-dimensional protein maps on anatomical models of the nephrogenic program. Single cell RNA sequencing identified novel progenitor states which were spatially mapped to the nephron anatomy enabling the generation of functional gene-networks predicting interactions within and between nephron cell-types. Network mining identified known developmental disease genes and predicts new targets of interest. The spatially resolved nephrogenic program made available through the Human Nephrogenesis Atlas (https://sckidney.flatironinstitute.org/) will facilitate an understanding of kidney development and disease, and enhance efforts to generate new kidney structures.

## Introduction

Birth defects across all organ systems often manifest as gross anatomical and cellular changes. These are difficult to link to specific developmental events or cell-types. While single-cell-omic approaches now facilitate the dissection and cataloguing of adult organs into their cellular components at an RNA and chromatin level (Macosko et al., 2015; Han et al., 2018; Schaum et al., 2018), performing equivalent analyses on developmental and disease processes is challenging. Tools that extrapolate developmental trajectories from fluid cell profiles differentiating from one state to the next tend to underrepresent the number of endpoints (precursor or terminally differentiated cell-types), erroneously group cells onto the same trajectory, and lose key spatial and dynamic information critical to a mechanistic understanding of the developmental program

The importance of understanding human developmental programs - i.e. the sequential series of differentiation steps by which progenitors and precursors give rise to their mature counterparts - is well illustrated by malformations of the genitourinary system (**Figure 1A**). Around 20-30% of all neonatal anomalies map to the genitourinary system accounting for 50% of pediatric end-stage kidney disease (Schedl, 2007; Hildebrandt, 2010). Once formed, the kidney controls homeostasis through blood pressure regulation and filtration, where metabolites are excreted and solutes recovered, predominantly through the actions of highly specialized cell types within distinct segments of the nephron (Lee et al., 2015). All nephron cell types are thought to arise from an equipotential nephron progenitor pool established at the onset of kidney development (McMahon, 2016). Recent single cell sequencing and spatial mapping studies indicate there are at least 23 cell-types within the nephrons of the adult male and female mouse kidney (Ransick et al., 2019). Considerable cell diversity has been reported for the human kidney (Lake et al., 2019) and it is likely this will expand as the depth and resolution of adult human studies increases.

**Figure 1.**
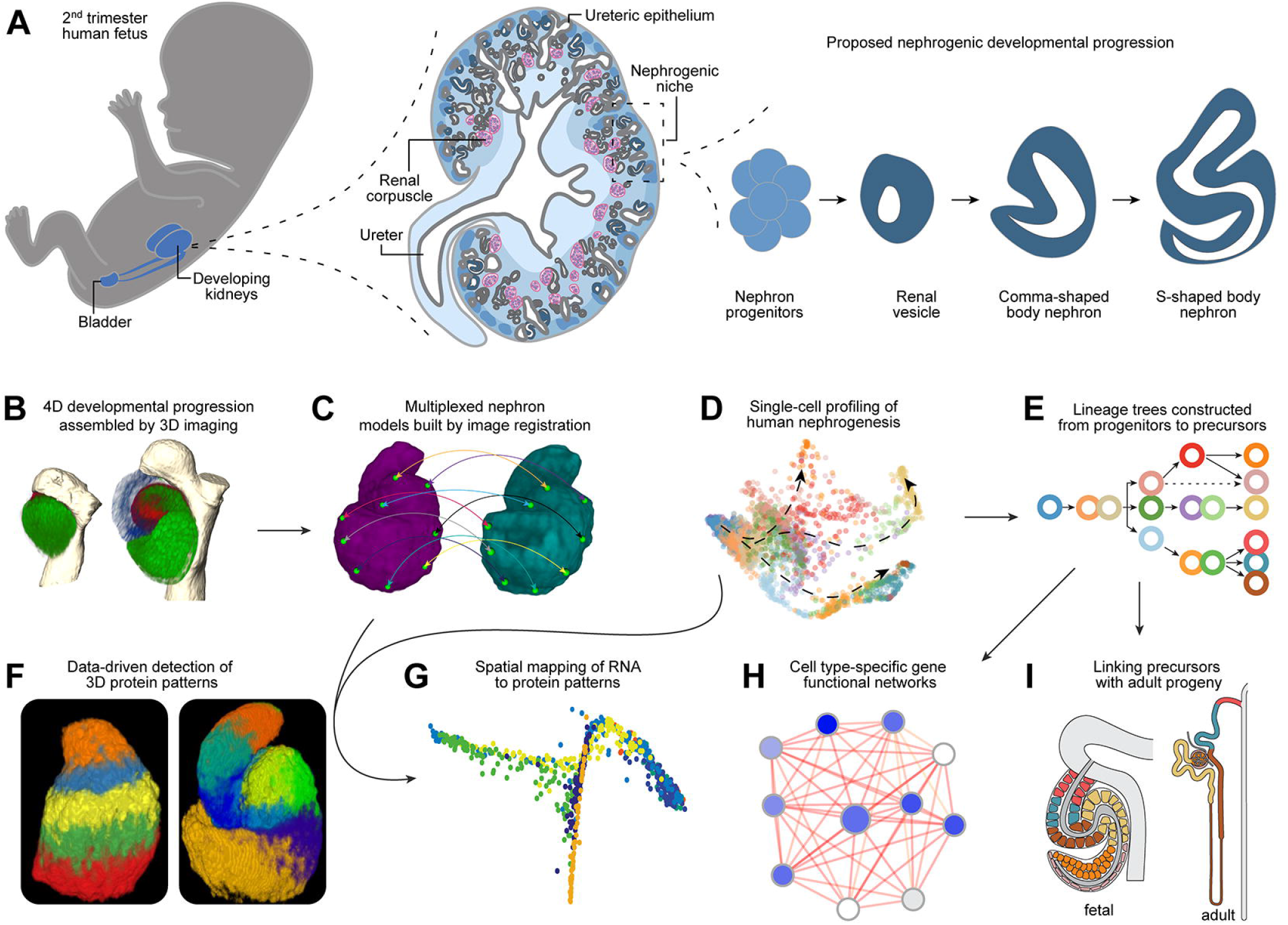
Assembly of the Human Nephrogenesis Atlas. Schematics and data illustrating: (A) Basic principles of human nephrogenesis. Schematic depicts genitourinary system in human fetus (left), a cross section of a developing human kidney (middle), and the proposed nephrogenic program (right). Dashed lines indicate magnified portions shown in a left to right direction. Shades of blue from lightest to darkest in kidney cross section indicate inner medulla, outer medulla, cortex, nephron progenitors, developing nephrons. Ureteric epithelium is shown in grey and maturing nephrons beyond the S-shaped body stage in dark grey. Renal corpuscles as indicated (pink/blue). (B) Visualization of immunofluorescently labelled nephrons in 3D. (C) Image registration of nephron protein data into multiplexed models. (D) Single cell RNA profiling of human nephrogenesis. (E) Establishing a lineage tree for nephrogenesis. (F) Computational approaches to predict protein patterns in 3D space. (G) Spatial mapping of RNA to protein models. (H) Building cell-type specific functional gene networks. (I) Linking precursors with their mature progeny.

The nephrogenic program has been well studied in mammalian model systems, principally the mouse (McMahon, 2016; **Figure 1A**). Mesenchymal progenitor cells for the nephron and interstitial cell lineages overlie epithelial progenitors for kidney’s collecting system, generating a highly interactive mobile nephrogenic niche that drives kidney assembly. Wnt9b signaling to mesenchymal nephron progenitor cells triggers commitment to the nephrogenic program (Carroll et al., 2005). Nephron progenitors cluster into pretubular aggregates, then epithelialize to form cystic renal vesicles, which transition through morphologically distinct stages (comma and S-shaped bodies), interconnecting with the epithelium of the developing collecting system, to establish the kidney’s fluid transporting epithelial network. Though precursor-product relationships are not well understood, growth and morphogenesis of the nephron anlagen is coupled with the emergence of dynamic profiles of gene expression along a proximal-distal axis of epithelial specialization that prefigures and determines the molecular and cellular organization of the adult nephron (Stark et al., 1994; Carroll et al., 2005; Kobayashi et al., 2008; Georgas et al., 2009; Karner et al., 2011; Lindström et al., 2018b, 2018a).

In previous efforts to resolve the diversity of nephron precursors, we and others have applied single cell RNA sequencing (scRNA-seq) to human nephrogenesis (Lindström et al., 2018a; Menon et al., 2018; Hochane et al., 2019; Tran et al., 2019). When cell clusters are compared with *in situ* and immunolocalization studies (Lindström et al., 2018b, 2018a), distinct cell clusters are likely an amalgamation of multiple related precursor types. Delineating the stepwise program of differentiation by which nephron progenitors generate nephrons will facilitate our understanding of disease origins. Further, the application of insight to stem cell systems will enable both the modelling of kidney disease in vitro, and de novo generation of kidney cell types and kidney structures.

To establish the spatial organization, diversity, and gene expression profiles of human nephron precursors we adopted and developed several approaches that are broadly applicable to other developmental systems. The data are brought together as a cohesive map of protein distribution, gene expression, and gene networks within an interactive database (https://sckidney.flatironinstitute.org/) that will facilitate study of the human kidney.

## Results

An overview of the workflow and projected goals underlying the generation of an atlas of human nephrogenesis is provided in Figure 1. Confocal imaging and image segmentation of the human nephrogenic zone with distinct antibody sets enabled the assembly of a 4-dimensional map of human nephrogenesis (**Figure 1A, B**). Image registration with these data was then used to generate multiplexed nephron models (**Figure 1C**). Extensive scRNA-seq analysis was performed on the nephrogenic zone (**Figure 1D**), transitional cell types verified *in vivo* (**Figure 1E**), and multiplex protein patterns (**Figure 1F**) were employed for relational mapping of transcriptomic data to a computational model of a developing nephron (**Figure 1G**). Cell typespecific functional gene-network models were combined with machine learning to examine genes associated with congenital abnormalities (CAKUT) and predict potential disease associated partners (**Figure 1H**). Finally, predictive precursor-product relationships were framed from the analysis of developmental and adult scRNA-seq datasets (**Figure 1I**). The following sections provide a detailed insight into the approaches, and the collection, analysis and interpretation of data, at each step.

## Visualizing progressive development of the human nephron

To visualize how mammalian nephrons form, we performed whole-mount immunolabeling of mouse and human nephrogenic niches with JAG1 antibodies as a reference landmark across nephrons, experiments and samples, in combination with antibodies recognizing eighteen other informative regional protein markers, incorporating DAPI labelling to resolve nuclei. Selection of antibody targets was designed to maximize detection of cell diversity in conjunction with nephron patterning and morphogenesis and to link development to disease given the known developmental roles of many of these proteins in mouse and human kidney studies. Two hundred and fifty-one human and one hundred and seventy-seven mouse nephron precursors were captured at high resolution by confocal imaging, digitally isolated from image stacks, and categorized into developmental stages based on gradual morphological changes, pattern, and size. Together, these data resolved in 3-D detail the developmental progression for human and mouse nephrogenesis to the S-shaped body stage (**Figure 2A; Supplementary Figure 1A-C**).

**Figure 2.**
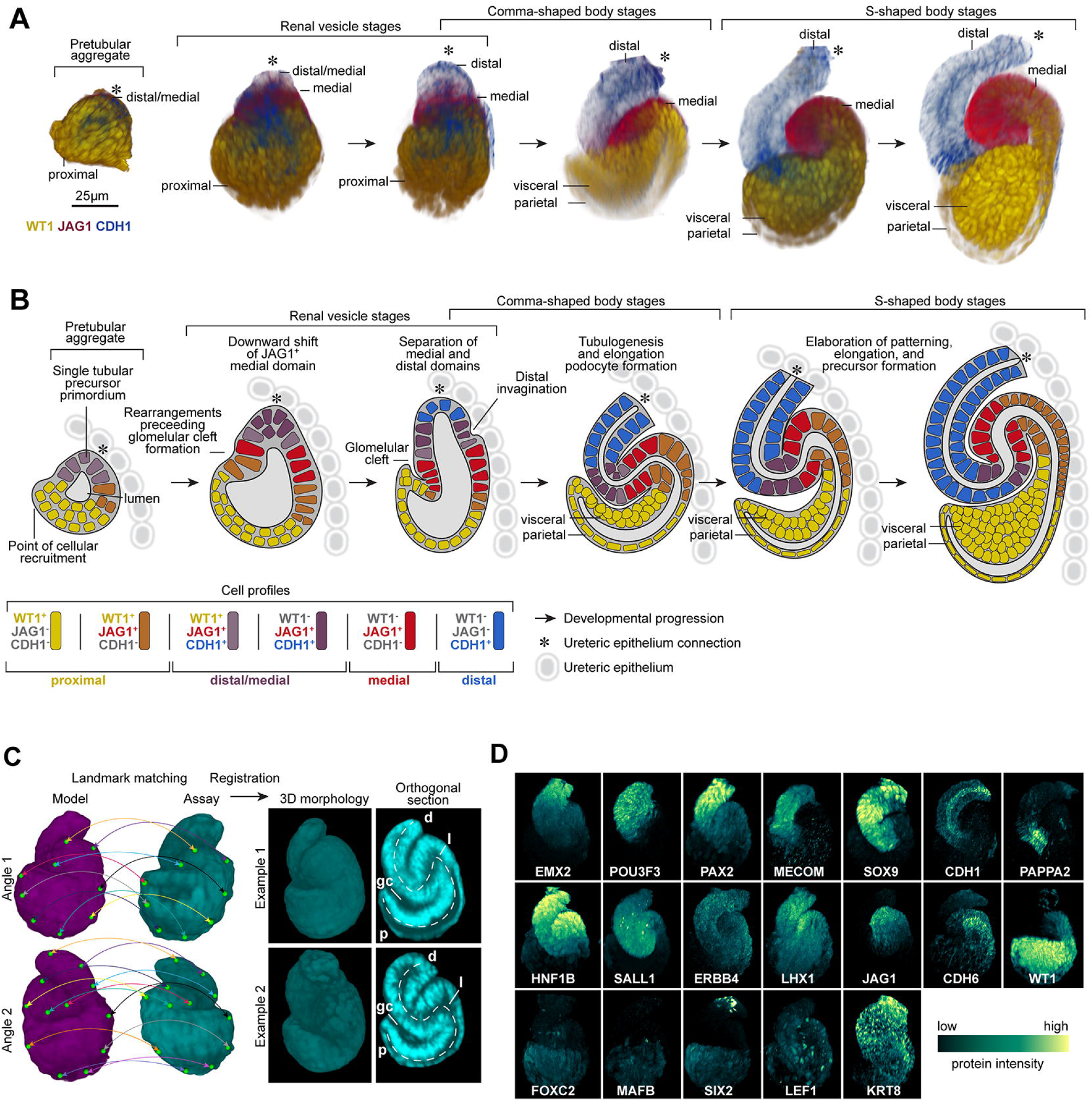
The developmental progression of human nephrogenesis. (A) Immunofluorescently labelled human nephrons rendered in 3D shown from most immature stages (left) to most mature stages of S-shaped body nephrons (right). (B) Schematic for nephrogenesis for an equivalent range of nephrons as shown in A, displaying separation of precursor identities over time. Immunofluorescent labels, features, and developmental stages as indicated on fields. Key defines cell profiles and markers. (C) Strategy for image registration and examples of registered 3D S-shaped body nephrons. A multi-stage registration process that utilized landmarks (green marks and colored lines linking matching landmarks), a general affine transform, and an elastic (ANTs) transform. Successfully registered examples of two S-shaped body nephrons shown at as orthogonal sections through the middle at two angles. Dotted lines indicate proximal distal axes. p: proximal. d: distal. gc: glomerular cleft. l: lumen. (D) Average localization patterns for proteins based on registered nephrons. Immunofluorescent labels as indicated on fields.

Nephron development initiates within a pretubular aggregate of tightly packed nephron progenitors. The epithelial transition visualized by CDH1 progresses from distal-to-proximal generating an apical epithelial surface shaped like an inverted funnel opening into a space that later will incorporate the kidney’s glomerular filter. Epithelialization begins in a distal JAG1+/ CDH1+ domain. In the epithelial renal vesicle, the distal half resolves into two domains: the most distal cells are now CDH1^+^/JAG1^-^, while more medial cells are CDH1^+^/JAG1^+^ (**Figure 2A-B**). An additional CDH1^-^/JAG1^+^ domain emerges medially during the transition to comma- and S-shaped bodies (**Figure 2A-B**). During the transition from renal vesicle to comma-shape body, inward folding between the medial and distal segments results in a distal orientated bend in the tube, while at the proximal end, epithelial movements establish the glomerular cleft. Vascular endothelial cells invading the cleft subsequently establish the vascular supply for the renal filter.

A schematic model for human nephrogenesis based on the cumulative view of these data (**Figure 2B**) is consistent with a gradual distal-to-proximal progression in nephron formation (Lindström et al., 2018a). Development of the mouse nephron over a similar developmental time course was similar (**Supplementary figure 1B-C**) though human nephrons were approximately twice the size of their mouse counterparts with two-fold greater cellularity (**Supplementary figure 1D**), consistent at an individual marker/domain level where CDH1^+^/Cdh1^+^ and JAG1^+^/Jag1^+^ cell numbers matched this two-fold difference and displayed similar levels of cell variation (**Supplementary figure 1E, F**); indicative of linear scaling of domains. The 3-D morphologies and protein patterns were consistently sharper in the human nephron across > twenty-five proteins. The size difference and the slower developmental pace of human nephrogenesis, roughly eight times slower to S-shaped body stage than the mouse kidney (Lindström et al., 2018c), likely contribute to the enhanced resolution of human nephrogenesis in our data.

### Image registration predicts stereotyped anatomies

Nephron formation within and across mammalian species follows a highly consistent early developmental progression at different stages of kidney development though there are clear differences in the final pattern and cell composition depending on the time of nephron formation (Ransick et al., 2019). To address whether nephrogenesis is stereotypical, we selected digitally isolated renal vesicles and S-shaped bodies from mammalian kidneys of distinct developmental ages, and performed image registration using an affine transform and elastic registration, to determine whether similar structures could be brought into the same voxel space, with minimal distortions to morphology or patterning. Nephron anatomies and fine anatomical details such as the lumen space and the forming glomerular cleft within the S-shaped body nephron were preserved, following registration (**Figure 2C**), thus indicating the low morphological variation in the nephrogenic program. However, S-shaped body nephrons displayed either a left-handed or right-handed spiral, indicating chirality depending on the orientation to the adjacent ureteric epithelium (**Supplementary figure 1G**). As expected, proteins were distributed in distinct distal-to-proximal patterns (**Figure 2D**). The multiplex nephron models for both renal vesicles and S-shaped body nephrons in themselves provide high-resolution views to the exact positioning of proteins with known roles in genetically linked developmental renal syndromes such as CAKUT (congenital anomalies of the kidney and urinary tract); for example: POU3F3, PAX2, HNF1B, SALL1, LHX1, and WT1. In addition, each distinct contribution enabled the creation of a high-resolution map of protein-defined, cellular diversity (**Supplementary figure 1H-I**), which is predicted to correlate with underlying differences in gene expression for each cognate gene within distinct cell populations. Importantly, the ability to co-register different nephron precursors within a kidney, and between kidney samples at different ages (13 to 17 weeks), shows mammalian nephrogenesis is a stereotypical process following a tightly controlled molecular and cellular program.

### Single-cell transcriptomes identified for human nephrogenesis

As the next step to the integration of scRNA-seq datasets to the spatial maps, we performed independent scRNA-seq analyses on two week-14 kidneys obtaining 24,157 cells displaying a median of 2,644 genes/cell (**Supplementary figure 2A**). Cells with a gene/cell count greater than 3,000 (8,316 cells) were analysed and non-cycling cells of the nephrogenic lineage (2893 cells) sub-setted. This high-quality nephrogenic cell set was evaluated further through reiterative unbiased clustering and gene enrichment analyses (see **STAR Methods** for details and **Supplementary figure 2A-J; Supplementary table 1**).

Eighteen distinct transcriptional states were identified using SEURAT (Satija et al., 2015) and by examining differential expression of key marker genes (**Figure 3A; Supplementary table 2**). To establish the range of sampled transcriptomes in clusters, we analyzed expression profiles for genes that decrease slowly after the progenitor state (O’Brien et al., 2016: *SIX1*), genes that are transiently upregulated in pretubular aggregates through to S-shaped body nephrons and then down-regulated (Lindström et al., 2018d, 2018b; Tran et al., 2019: *CAPN6, CRYM, OLFM3, JAG1;* **Supplementary figure 3A**), and genes that are first activated in S-shaped body nephrons (Lindström et al., 2018d, 2018b; Tran et al., 2019: *NPHS1, CDH6, HNF4A, MECOM, GATA3, PAPPA2;* **Supplementary figure 3B**). While false negatives can challenge interpretations of individual cells in scRNA-seq data, at a cluster level each of these genes were detected robustly. The sampled cells ranged from nephron progenitors to S-shaped body nephron cells. Six clusters were identified as S-shaped body nephron cells (**Supplementary figure 3C**) that displayed deep transcriptional differences (**Supplementary figure 3D**). The 18 clusters expressed on average 12,900 genes - likely a good representation of each cell type’s full transcriptome. Importantly, three-fold more clusters were detected for the S-shaped body compared to previously reported data (Lindström et al., 2018a; Menon et al., 2018; Hochane et al., 2019; Tran et al., 2019) and the depth and breadth of the resulting transcriptomic data enabled subtle gene expression changes to be visualized within scarce transitioning cell-states.

**Figure 3.**
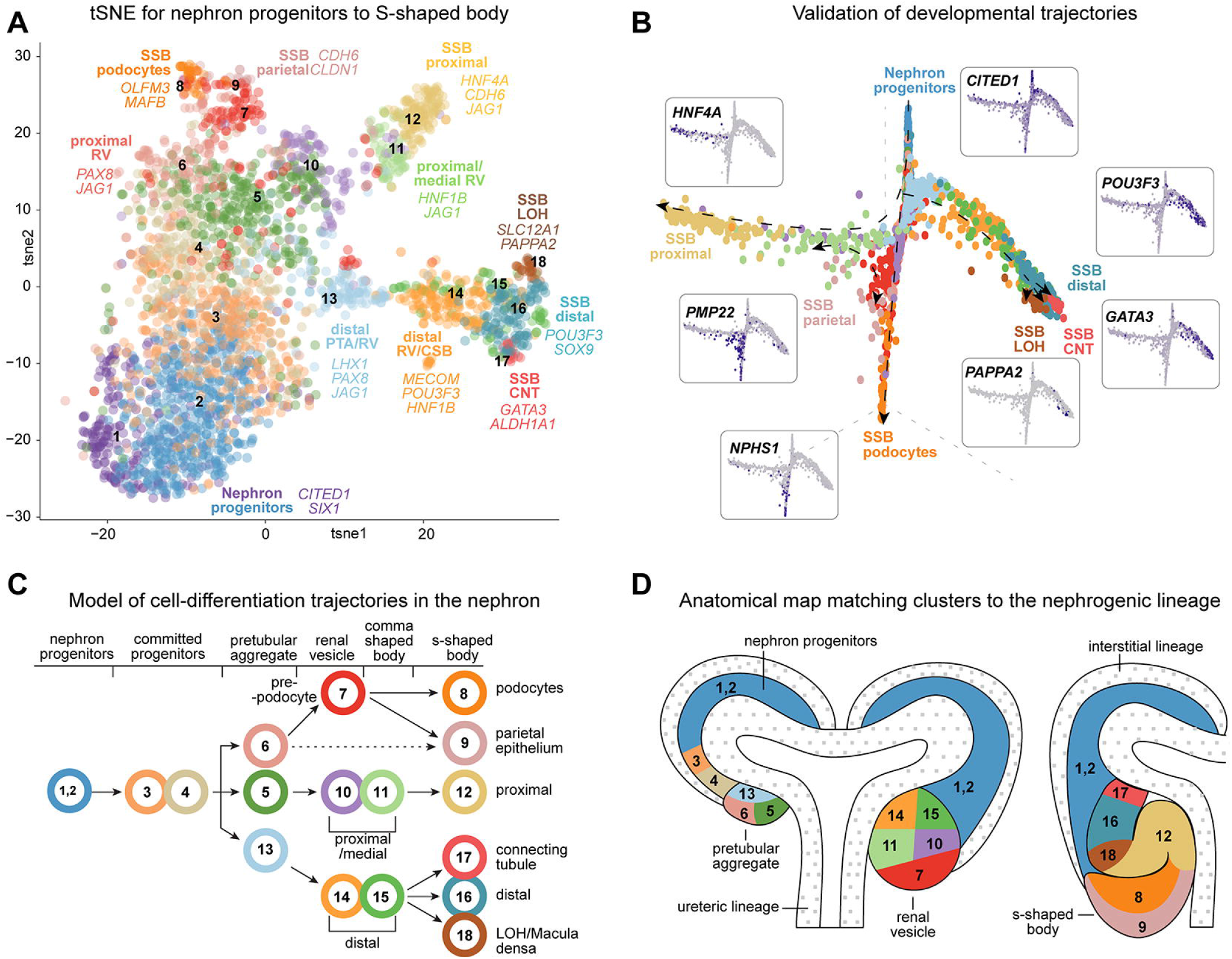
Single cell analyses and spatial mapping resolve precursor transcriptomes. (A) tSNE plot of 18 cell clusters of cells from human nephrogenic lineage with cell-types and marker genes indicated. (B) Graph-based dimensionality reduction plots of all cell transcriptomes (clusters 1-18) color coded according to cell clustering in Figure 3A. Marker genes for each endpoint shown as inserts. (C, D) Lineage tree model for nephrogenesis placing cell transcriptomes in clusters 1-18 into a tree formation and into an anatomical schematic for the nephrogenic niche. Colors and numbers correspond to cluster colors in Figure 3A. Arrows indicate direction of differentiation. Dotted arrow indicates uncertain relationship. Stages and precursor regions or identities as annotated. White areas with small grey dots in D mark the ureteric epithelium and the interstitium which did not form part of these analyses.

### Resolving changing expression profiles across time and anatomy

To resolve cell states between nephron progenitors and the S-shaped body cell populations we used graph-based dimensionality reduction (DRGraph) (Menon et al., 2018). Three main trajectories were evident, starting from nephron progenitors (clusters 1 and 2) and ending in cells displaying markers of distal (cluster 16, 17, 18), proximal (cluster 12), podocyte (cluster 8), and parietal epithelium (cluster 9)(**Figure 3B; Supplementary figure 4A-D**). These cluster relationships agreed with correlation analyses of all variable genes (12995) across the clusters (**Supplementary figure 4E**) which similarly grouped clusters into distal, proximal, and podocyte/parietal categories. Thus, DRGraph trajectories, correlation analyses of all variable genes across all cells, and SWNE projections (**Supplementary figure 2J**) suggested a consistent developmental ordering of the clusters.

To annotate clusters with developmental and anatomical terms we used RNAscope to simultaneously visualize the expression of multiple genes. The genes have predicted overlapping but dynamically different expression profiles across anatomically distinct developmental stages facilitating the assignment into precise anatomical bins (**Supplementary figure 5A-D;** the predicted expression domains in **Supplementary figure 5E-J**). *COL4A4/TMEM72/SLC39A8* were used to annotate clusters as distal patterns emerged, while *TMEM72/SLC39A8/LAMP5* were used to annotate clusters belonging to the proximal domain as it emerged from the shared distal population in the pretubular aggregate and renal vesicle. *SLC39A8/LAMP5/HNF4A* facilitated resolving cell identities of proximal cells in the S-shaped body, with very rare *SLC39A8/HNF4A* and common *HNF4A/LAMP5* double positive cells. *SLC39A8* straddles the border between distal and medial S-shaped body domains. *COL4A4/HNF4A/EFNB2* identified cell fates formed during the differentiation of distal, proximal and podocyte fates, respectively (**Supplementary figure 5A-J**). Clusters annotated with anatomical and developmental terms thus form an anatomical model for the 18 transcriptomes that predicts hierarchical relationships amongst the cell types and potential relationships to the adult nephron (see later data and discussion; **Figure 3C-D**). Collectively, these data have been incorporated into a spatially resolved and searchable Human Nephrogenesis Atlas (NephMap - https://sckidney.flatironinstitute.org/).

### Unsupervised protein clustering predicts novel precursor populations

Twelve distinct transcriptomes characterized the progression of nephrogenesis from renal vesicles to S-shaped bodies (**Figure 3C-D**). To determine if protein combinations in the nephron models could be formalized into distinct combinations of protein abundance corresponding to precursor populations, we performed k-means clustering treating each voxel within the multiplexed 3D models as an n-dimensional vector in which each dimension corresponds to the abundance of a single protein. For renal vesicles we based clustering on the localization of 18 proteins, while for the S-shaped body nephrons we performed clustering on 19 proteins (**Figure 4A**). In renal vesicles, patterns formed increasing numbers of parallel layers along the distal-proximal axis as increasing k-values were used as the input, thus indicating proteins were distributed in gradients along the proximal distal axis (**Supplementary figure 6A**). By the S-shaped body stage however, eight patterns were predicted in separate distinct regions along the proximal-distal axis. These patterns were stable across increasing k-values, suggesting each consisted of a unique protein combination with defined boundaries (**Figure 4A; Supplementary figure 6B**). The S-shaped body patterns agree with previously postulated precursor domains (Lindström et al., 2018b). The protein patterns and abundance of proteins within each pattern support the model that the tubular portion of the nephron, excluding podocytes and parietal epithelial cells of the renal filter, forms from a single precursor domain, which separates as development progresses into distal and medial domains (**Figure 4B-C**). For example, initially CDH1 and JAG1 overlap, then separate into distal and medial domains (**Figure 4B-C; Supplementary figure 6E; Figure 2B**). However, other proteins such as EMX2 and HNF1B remain co-localized (**Figure 4B-C; Supplementary figure 6E**).

**Figure 4.**
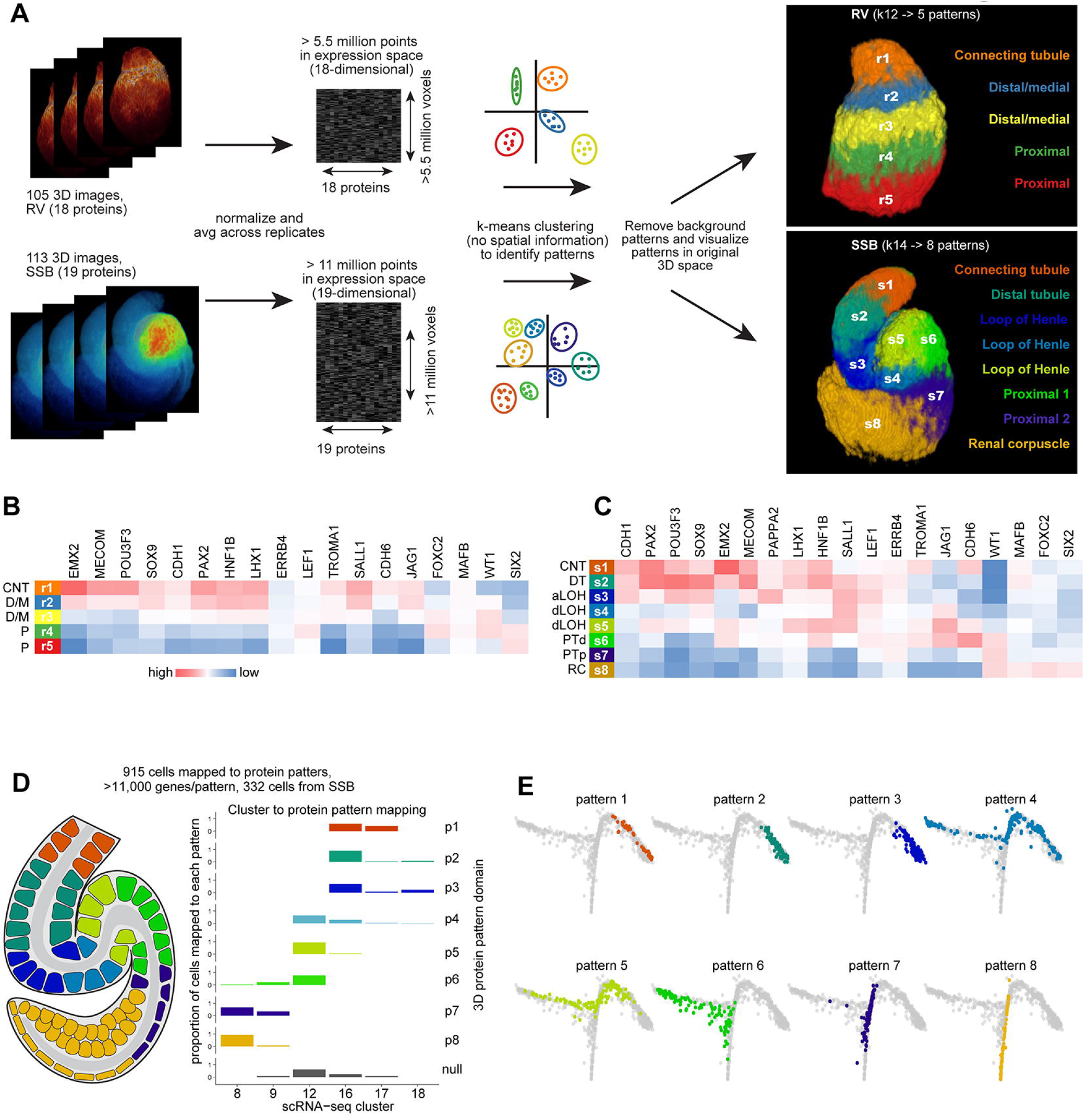
Unsupervised clustering approaches predict precursor populations. (A) Workflow of computational approach used to apply k-means clustering to 3D multidimensional voxel-based data and resulting predictions of combinatorial protein patterns. (B, C) Protein abundance across predicted protein patterns in renal vesicles and S-shaped body nephrons. (D) Schematic of mapping of cells to 8 protein patterns in the S-shaped body nephron and proportion of cells of the selected clusters mapping to each protein pattern. (E) Graphbased dimensionality reduction plots of all cell transcriptomes (clusters 1-18) showing cells spatially mapped to S-shaped body patterns. Colors in E correspond to protein patterns as shown in D. CNT: connecting tubule. D/M: distal/medial. P: Proximal. DT: distal tubule. aLOH: analage of loop of Henle. dLOH: putative analage of descending loop of Henle. PTd: analage of proximal tubule. PTp: analage of proximal tubule and parietal epithelium. RC: renal corpuscle.

### Three-dimensional spatial mapping of single-cell transcriptomes

Recognizing mRNA levels will only approximate levels of encoded protein products, we mapped cell expression profiles to the 3D protein models based on gene-to-protein relationships (**Figure 4D**). Three hundred and thirty-two S-shaped body cells mapped to the 8 patterns with >11,000 genes detected per pattern. Differentiation trajectories were predicted with DRGraph for cells transitioning from the pretubular aggregate to S-shape body stage and cell groupings were mapped onto these trajectories (**Figure 4E; Supplementary figure 6F**).

In reconciling the gene expression profiles generated by spatial mapping to S-shape body nephrons with those from conventional clustering algorithms (**Supplementary figure 7A vs. Supplementary figure 3D**), several genes highlighted how different views are reached by these methods. *SLC39A8*, encoding a transmembrane zinc transporter exhibiting preweaning lethality in mouse mutants (International Mouse Phenotyping Consortium), was by conventional gene enrichment analysis differentially expressed in clusters 13, 14, 15, (**Supplementary table 2**) but not cluster 16 (**Figure 3D and Figure 4A, D**). However, differential gene expression based on spatially mapped cells showed *SLC39A8* to be expressed in a distal subdomain of the S-shaped body nephron equivalent to pattern 2, and confirmed *in vivo* (**Supplementary figure 7A-, B**). The increased temporal resolution provided by spatial mapping of transcriptomes therefore accurately identified *SLC39A8* as a differentially expressed gene marking a previously uncharacterized region of the S-shaped body nephron. Likewise, *LAMP5*, a lysosome-associated membrane glycoprotein, which in mutant mice display neurological defects (Tiveron et al., 2016) was predicted by transcriptomic analyses to be enriched in clusters 7, 9, and 11 but not in S-shaped body clusters. However, expression was predicted by spatial mapping and observed *in vivo* in a narrow region of the proximal S-shaped body, matching pattern 6, again suggesting spatial mapping can improve resolution to gene expression patterns and identify subdomains of cell-populations (**Supplementary figure 7C**). A most striking example showing complementary strengths of applying spatial transcriptomics is *HNF4A*, a transcription factor required for proximal tubule development and altered in CAKUT (Marable et al., 2018). It was predicted to overlap with *LAMP5* in the proximal S-shaped body in pattern 6 and this prediction was borne out despite their divergent expression at earlier stages of nephrogenesis (**Supplementary figure 7D**). This maps HNF4A to a very small domain of the S-shaped body (discussed further in Figure 6). In summary, mapping of RNA profiles to protein domains is feasible. Further, integrating multiple modalities in the relational mapping of RNA profiles gives detailed molecular insight into cell diversification underlying nephron differentiation.

### Beyond the single gene: gene networks in development and disease

To build a richer view of gene expression environments and pinpoint cell-disease/dysgenesis relationships, we applied a regularized Bayesian machine learning approach to build cell typespecific functional gene networks combining the scRNA-seq data with ~1500 publicly available human genomic datasets (Greene et al., 2015; Wong et al., 2018) (**Figure 5A**; https://sckidney.flatironinstitute.org/genenetwork). To validate this approach, we first explored genes with known expression, function and interacting partners. *JAG1*, which is active in proximal nephron development (Liu et al., 2013), was identified as being tightly connected to Notch pathway components *NOTCH1*, *NOTCH2*, *NOTCH3*, *HES1*, and *ADAM9* in the proximal/medial renal vesicle network where *JAG1* was strongly expressed (**Figure 5B**) but as expected not in the distal most connecting tubule region where *JAG1* expression was absent (**Supplementary figure 8A**).

**Figure 5.**
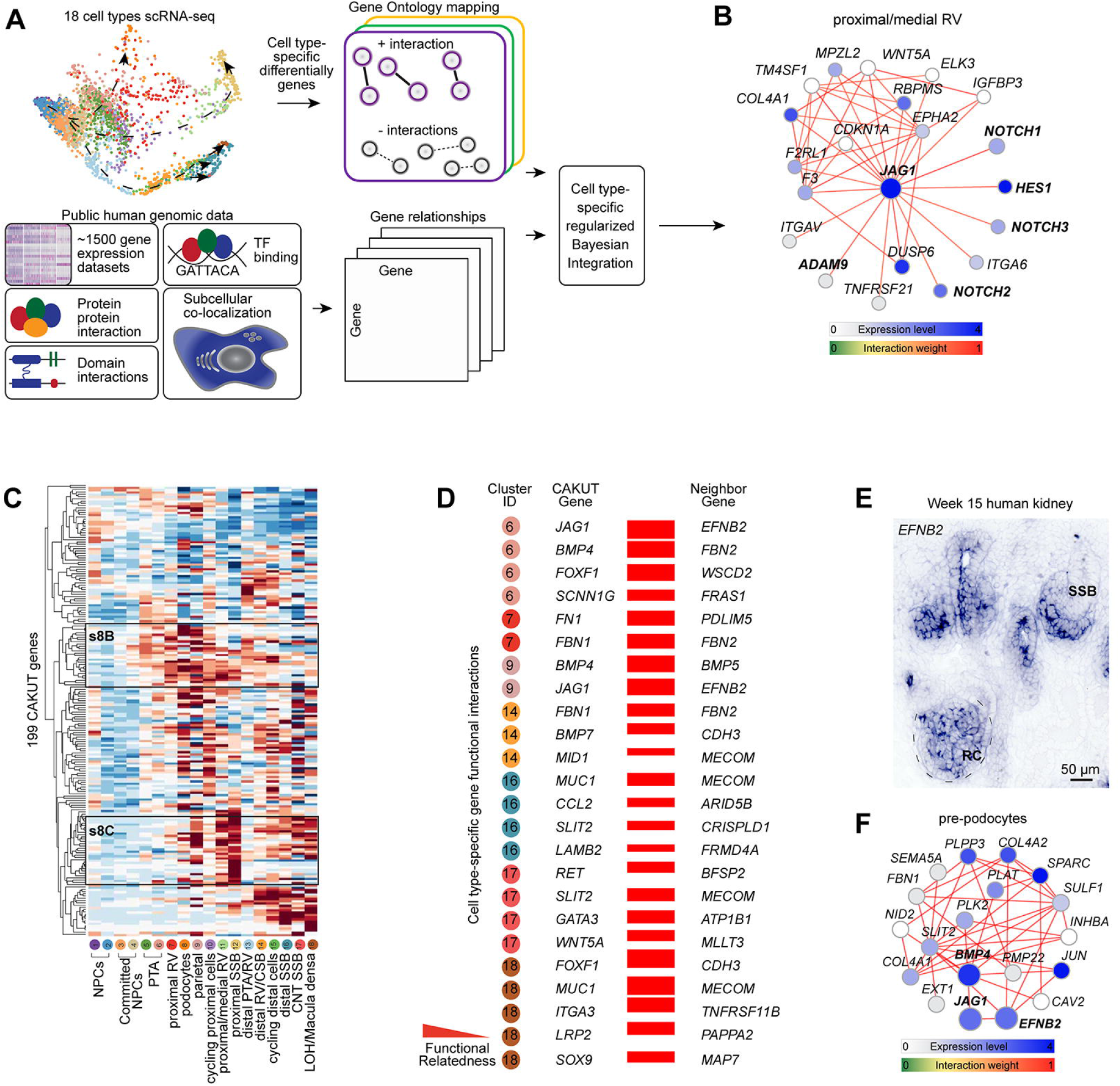
Cell-type specific functional gene networks in development and disease. (A) Schematic for building cell-type specific functional gene networks. (B) Functional gene network for JAG1 (in bold) in the proximal/medial renal vesicle (cluster 11). (C) Hierarchical clustering of 199 CAKUT genes across cluster identities 1-18. High/Low as indicated. S8B and s8C annotations indicate regions highlighted in Supplementary figure 8B, C. (D) CAKUT genes and their strongest neighbor genes. (E) In situ hybridization for *EFNB2* on human fetal kidney. (F) Example of triple gene network for *EFNB2, JAG1*, and *BMP4* – in bold. Scales and stains as indicated. Expression levels and edge weights as indicated by scales on panels. PTA: pretubular aggregate. RV: renal vesicle. CSB: comma-shaped body nephron. SSB: S-shaped body nephron. CNT: connecting tubule. LOH/MD: loop of Henle/Macula densa precursor. RC: renal corpuscle. Gene network thresholds set for visualization purposes to ensure network simplicity.

To focus on dysgenesis and disease, 272 genes published and linked to kidney disease and CAKUT (Hildebrandt, 2010; Vivante et al., 2014) were hierarchically clustered based on their average gene expression in the Human Nephrogenesis Atlas. Of these, 199 genes were detected in distinct expression groups (**Figure 5C**). Proximal epithelial cells abutting the podocytes showed strong expression of the lipid and solute transporter proteins, *LRP2, SLC7A9, SLC3A1, SLC5A2* components of adult proximal tubule cells; (**Supplementary figure 8B**). *PAX8* and *WNT4* defined early differentiation and epithelialization stages, and other gene sets multiple pathway components; for example, (e.g., proximal biased expression of the Notch pathway genes *JAG1, NOTCH2* and *LFNG*) (**Supplementary figure 8C**), consistent with published mouse and human studies (Chen and Al-Awqati, 2005; Cheng et al., 2007; Chung et al., 2016; Lindström et al., 2018b). Genes linked to a range of renal disease categories (nephrotic syndrome, renal tubular acidosis, kidney stone, CAKUT) have previously been mapped to adult mouse cell transcriptomes (Park et al., 2018). Analysis of the 20 CAKUT genes evaluated by Park et al. highlights the complementary insight from comparisons to developmental data. As examples, *EYA1* and *SALL1* are required for nephron progenitor selfrenewal (Basta et al., 2014; Xu et al., 2014), which is consistent with our expression data, while in the adult expression data, *EYA1* and *SALL1* map to podocytes and unknown cell-types that may not relate to CAKUT origins (Park et al., 2018; Ransick et al., 2019) (**Supplementary figure 8D**). In contrast, *JAG1* is known to have multiple functions in the kidney. JAG1 is required for patterning of the nephron (Cheng et al., 2007; Liu et al., 2013) and in the differentiation and maintenance of collecting duct cell-types in the adult kidney (Chen et al., 2017). Developmental and adult datasets identify each cell population (**Supplementary figure 8D**). Collectively, direct comparison of the 199 CAKUT genes shows a stronger association with relevant cell types within the developmental data, as expected for a developmental category of disease (**Supplementary figure 8E**).

To predict novel candidate genes for association studies to developmental disorders and CAKUT, we selected genes whose cell-type specific edge weights to CAKUT genes were significantly higher (Wilcoxon test, p-value <0.05) than non-CAKUT disease genes (OMIM database). Functional gene networks are a powerful tool to interrogate GWAS data and find genes not identifiable through conventional p-value cutoffs (Krishnan et al., 2016). We selected four GWAS studies covering 966,864 individuals with reduced glomerular filtration rate (GFR), a broad readout for abnormal kidney function (Hwang et al., 2007; Köttgen et al., 2009; Pattaro et al., 2016; Wuttke et al., 2019). One hundred and nine genes were strongly linked to CAKUT genes and detected through SNP analysis (p<0.05) in the GWAS studies. Seventeen of these were expressed in >10% cells in a distinct cluster with a functional connection weight score to the best CAKUT gene greater than 0.3 (candidate genes and the connected CAKUT genes were shown in **Figure 5D**).

Ephrin B2 (*EFNB2*) was identified in the pre-podocyte cluster (cluster 6) and likely parietal epithelium cluster (cluster 9) as a close neighbor to *JAG1*, a gene with strong links to Alagille syndrome and kidney abnormalities (Li et al., 1997; Spinner et al., 2001), and the regulation of angiogenesis (Benedito et al., 2009). *Efnb2* has previously been proposed to be involved in glomerular vascular assembly (Takahashi et al., 2001) and control of angiogenesis outside of the kidney (Wang et al., 2010). SNPs associated with *EFNB2* were identified in each GWAS study and we confirmed *EFNB2* expression in the developing renal corpuscle (**Figure 5E; Supplementary figure 8F**). *BMP4*, which was recently shown to be upregulated during early human renal corpuscle formation (Kim et al., 2019) was identified as differentially expressed in the pre-podocyte and parietal epithelium and also tightly connected to both *EFNB2* and *JAG1* in the corresponding networks. *Bmp4* is required for normal renal corpuscle formation in mice (Ueda et al., 2008). The combined *EFNB2, JAG1, BMP4* network in the pre-podocyte cells was enriched for GO terms indicating vascular development (**Figure 5F;** data not shown) suggesting the mixture of known and putative CAKUT genes regulate the formation of the renal corpuscle and its vascular architecture.

Expression of a second set of genes with strong connections to CAKUT - *MECOM*, *PAPPA2*, and *TNFRSF11B* - coincide in cluster 18 (**Figure 5D**). *MECOM* mutations were recently identified in patients with kidney malformations (Germeshausen et al., 2018), *Pappa2* is connected to salt-induced hypertension in rats (Cowley et al., 2016), and *TNFRSF11B*, also known as Osteoprotegerin, is linked to hypertension and chronic kidney disease (Bernardi et al., 2017) and was confirmed to be strongly expressed in the macula densa and also more broadly in the distal tubule (**Supplementary figure 8G, H**).

These examples highlight how novel candidate genes associated to developmental disorders and CAKUT can be identified by intersecting single-cell transcriptomics and functional gene networks.

### Exploring precursor-product relationships in the developing nephron

Our work identifies cell populations in the S-shaped body that expressing genes that in the adult nephron are restricted to distinct cell fates, thus making them putative precursors for adult nephron cell types. However, precursor-product relationships are poorly understood in the nephron, primarily due to the complexity of gene expression patterns where genes are activated multiple times in different nephron segments, thus preventing genetic lineage tracing experiments.

To explore precursor-product relationships, we performed correlation analyses between genes expressed by the six developmental S-shaped body cell-types and the orthologs expressed in the twenty-two resolved final fates found in the adult mouse. Adult podocytes, parietal epithelium, proximal tubule, distal portions of the loop of Henle, the distal tubule, and connecting tubule showed positive correlation to clusters 8, 9, 12, 18, 16, and 17, respectively (**Figure 6A** adult single cell data extracted from Ransick et al., 2019, which resolves the nephron into >22 transcriptonal signatures)). Adult cell signatures from the inner medulla and from specialized cells in the connecting tubule poorly correlated to any of the developmental transcriptomes.

**Figure 6.**
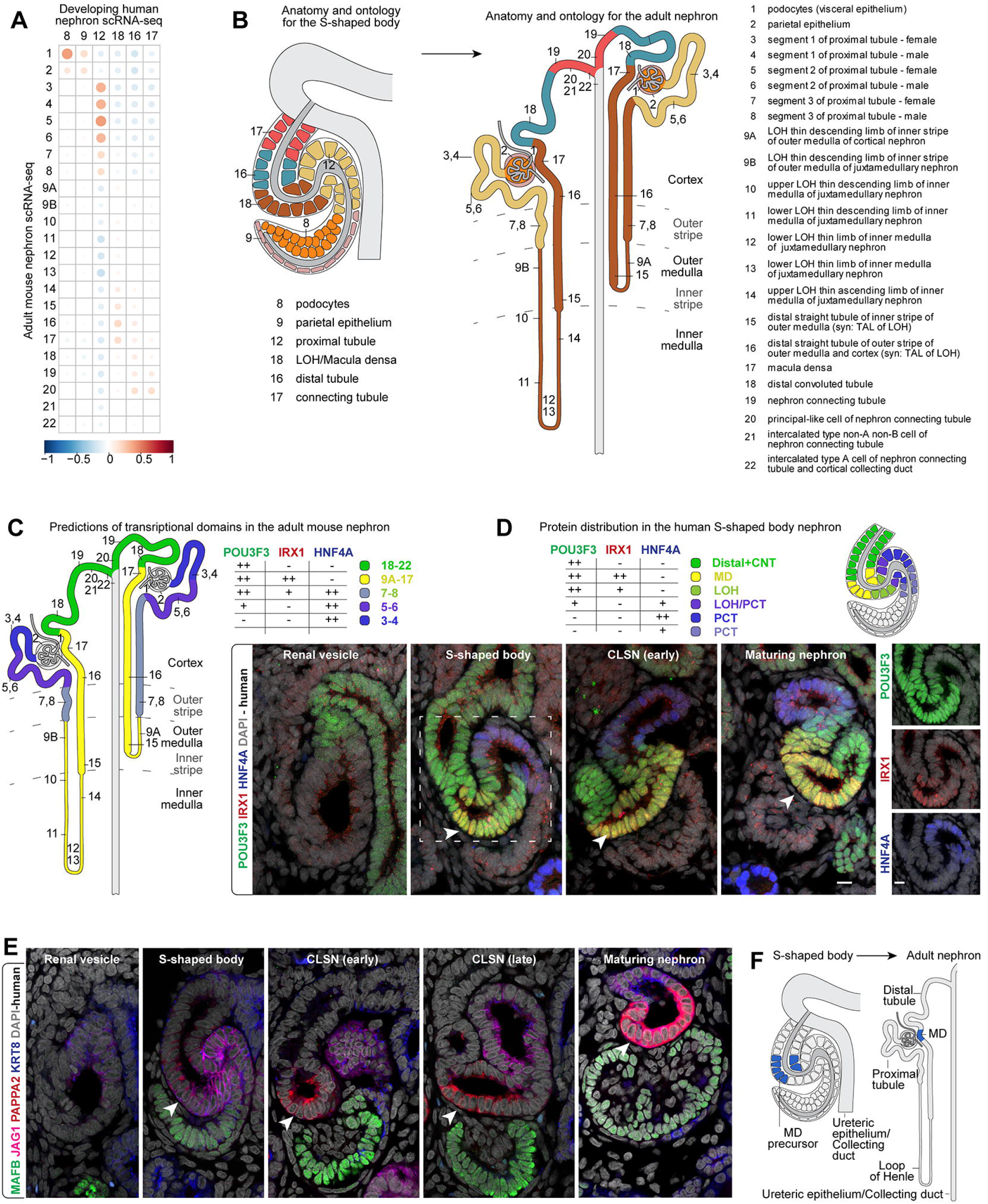
Linking precursors and progeny. (A) Correlation of gene expression between adult mouse clusters and human developmental clusters in S-shaoed body. (B) Proposed model for relationships between S-shaped body precursors and adult nephron cell-types based on correlation analyses. (C) Expression of transcription factors in the adult mouse nephron. (D) Immunofluorescent stains for transcription factors in human nephrons. White arrowheads indicate putative macula densa precursors. Schematics indicating the area each protein is detected in S-shaped body nephrons. Schematics also summarize data in Supplementary figure 11. (E-F) Immunofluorescent labelling of human nephrons during serial stages of development and schematic indicating relationship between macula densa precursors and mature derivatives. MD: macula densa. CLSN: capillary loop stage nephron. Labelling as indicated on fields.

Given that transcription factors impart and maintain cell fates we determined whether mouse orthologs of developmentally enriched human transcription factors that are expressed in the adult mouse nephron remain expressed in pattern resembling those in the developing nephron (**Supplementary figure 9A**). Many genes were expressed within domains that at a nephron structure-level corresponded to potentially relative matching positions along the adult proximal distal nephron axis. *TFAP2A* (human symbols given where orthologs have identical name between mouse and human) and *MECOM* were strongly expressed in distal S-shaped body segments and likewise restricted to distal segments in the adult nephron. *IRX1* and *IRX2* were strongly enriched in cluster 18 of the developing nephron and similarly enriched in the adult mouse macula densa. Other orthologs of cluster 18 enriched genes, e.g. *SIM1* and *HOXB7* showed expression both in the distal segments and the loop of Henle of the adult nephron, but were absent in proximal regions. Cluster 12 enriched genes, e.g. *HNF4A, HNF4G*, and *HNF1A* are in the adult nephron exclusively expressed in the proximal tubule indicating early programming of proximal tubule cell fates. Similarly, many transcription factors expressed by developing podocytes (*MAFB, WT1*, and *CREB3L2* - cluster 8), were detected in adult mouse podocytes. Cluster 9 showed few genes that were expressed in adult cells within the proximal portion of the nephron. Examining how orthologs of mouse transcription factors enriched within regions of the adult mouse nephron were expressed in the developing human nephron, a strong link was again found to cluster 18 of the S-shaped body and the loop of Henle and macula densa cells of the adult mouse kidney (**Supplementary figure 9B**). Consistent with an anatomical pre-positioning of progenitors, more proximal clusters 12 and 8 in the S-shaped body displayed strong expression of proximal tubule and podocyte enriched transcription factors, respectively.

Aside from podocytes which likely form in the S-shaped body, but have formally only been genetically traced from capillary loop stage nephrons to the adult (Eremina et al., 2002), we focused on two potential precursors populations, those for the macula densa (likely a subset within cluster 18) and proximal tubule (cluster 12). *Hnf4a* (the mouse ortholog of *HNF4A* enriched in cluster 12) has previously been suggested to mark mouse proximal tubule precursors and is functionally required for proximal convoluted tubule formation (Marable et al., 2018). Cluster 12 cells spatially mapped to a narrow region in the proximal S-shaped body, which we confirmed to be *HNF4A^+^* by *in situ* hybridization (**Figure 4D, E; Supplementary figure 7D**). We further resolved the HNF4A^+^ domain to a narrow cell-layer between proximal WT1^+^ and medial JAG1^+^ S-shaped body cells (**Supplementary figure 10A**). Thus, cluster 12 may incorporate a proximal tubule precursor population. Cluster 18 displayed highly enriched expression for *PAPPA2, IRX1, IRX2*, and *ERBB4* (**Supplementary figure 3D, Supplementary figure 9A**) indicative of a mixed cellular identity with characteristics of the distal loop of Henle, distal tubule and macula densa (Reggiani et al., 2007; Alarcón et al., 2008; Veikkolainen et al., 2012). The majority of cluster 18 cells mapped to protein pattern 3 (**Figure 4D**) and strikingly, PAPPA2 protein was detected in an equivalent region within the distal S-shaped body tubule adjacent to the forming renal corpuscle.

These data suggest PAPPA2^+^ cells comprise precursors to the macula densa. While lineage tracing is not possible in the human kidney, we tested whether PAPPA2^+^ cells overlap with macula densa cells using our recent zonal kidney scRNA-seq data where more mature stages of nephrons were sampled across the cortex and medulla of week 17 kidneys (Tran et al., 2019). A *PAPPA2^+^/NOS1^+^* population was identified (**Supplementary figure 10B-D**), *NOS1^+^* is highly enriched within the adult macula densa where NOS1-mediated NO signaling plays a key role in regulating renal blood flow (Mundel et al., 1992; Wilcox et al., 1992; Peti-Peterdi and Harris, 2010). Genes tightly correlating with *PAPPA2* in the week 17 data and that were differentially expressed in cluster 18 (S-shaped body data) included family-members of the Iroquois-class homeodomain transcription factors *IRX1, IRX2, IRX3, IRX5*, and *IRX6* (but not *IRX4*), whose expression closely followed *PAPPA2* (**Supplementary figure 10E, F**). To analyze cell-transcription factor expression in depth, we selected six transcription factors for immunolabelling (IRX1, TFAP2A, TFAP2B, EMX2, TFCP2L1, MECOM) and performed codetection with PAPPA2, and broader distal domain markers (POU3F3 and CDH1) in the S-shaped body (**Figure 6C-D; Supplementary figure 11A-F**). A small shared domain with overlap of all transcription factors exist and coincides with the narrow localization domain for PAPPA2. Most strikingly, careful temporal analysis of the PAPPA2 domain across developmental stages of nephrons showed how the PAPPA2^+^ domain emerges asymmetrically at the S-shaped body stage and persists in this configuration and in close contact with the developing renal corpuscle through to maturing stages where a PAPPA2+ macula densa could be identified anatomically (**Figure 6E**).

We therefore concluded that a small cell-population in the S-shaped body displays strong expression of markers identified in the adult macula densa, indicating this likely forms as a cell plaque in the S-shaped body nephron (model in **Figure 6F**).

## Discussion

We have established a developmental program for human nephrons beginning at multipotent nephron progenitors and ending with precursor populations of the S-shaped body nephron. Integrating spatial, transcriptomic and protein information provides a model for cell-relationships as nephron assembly progresses. The data describes a range of previously uncharacterized transient transcriptomic and protein states at each point of the differentiation trajectories, insights consolidated within the framework of the Human Nephrogenesis Atlas (https://sckidney.flatironinstitute.org/).

The data point to the emergence of cell-fates within a stereotypical program of epithelial morphogenesis which prefigures a progressive increase in cellular complexity. Consistent with lineage tracing studies in mouse nephron progenitor cells (Kobayashi et al., 2008), single cell analysis suggests there is a common precursor to distal and proximal epithelial tubular segments of the human nephron. Distinct proximal-distal programs correlate with proximal distal positioning within the renal vesicle and later structures, which is itself dependent on the time of recruitment of committed cells into the renal vesicle (Lindström et al., 2018a). How time of recruitment and position within the nephron anlagen ultimately determines cell identity remains to be determined though the integration of signaling networks is likely a key component (Lindström et al., 2018a).

The emerging view is that by the S-shaped body stage, many of the precursor populations begin to show some characteristics of transcription factor expression and function-associated gene expression profiles that pre-figure adult cell-types (**Supplementary figure 9**). This opens the opportunity to link precursors to their physiologically active progeny. One cell-type of particular significance is the macula densa, which regulates blood pressure through the reninangiotensin system (Peti-Peterdi and Harris, 2010). The macula densa is made up of around - 20-30 cells on the glomerular facing side of the distal nephron tubule that sense Na+ levels in the renal filtrate passing through the nephron, to modulate glomerular blood flow through local and systemic processes, controlling the rate of renal filtration (Peti-Peterdi and Harris, 2010). With 26% of the world’s population affected by hypertension (Kearney et al., 2005) understanding how cells form that control blood pressure, will facilitate the development of new strategies to manage hypertension.

Interestingly, cells in population 18 in the S-shaped body (**Figure 6A-E**) in the position of the second hair-pin bend along the proximal to distal trajectory of the convoluted nephron anlagen, shows several features of cells of the adult macula densa, which we map to a narrow asymmetric PAPPA2+ domain. As in the adult, these putative macula densa progenitors have a close association with the developing renal corpuscle suggesting an early functional anatomical association at the S-shape body stage, underpinning effective local signaling by macula densa cells regulating glomerular filtration rate in the adult kidney. Further, this interpretation suggests the conserved morphogenesis observed in the mammalian nephron forming program reflects, at least in part the early generation of cellular relationships essential to critical physiological regulation in the functioning kidney. Further, given the size of this cell population in the S-shaped body, and that of adjacent podocytes, relative to their adult cell numbers, there may be little further replication of these cells beyond the S-shape body, stabilizing their interactions, and local assembly of renal filtration systems. This model predicts a large component on nephron diversity (all PT segments, and the ascending and descending loops of Henle, emerge from a progenitor pool within the proximally adjacent populations (**Figure 6B, F**) of the S-shaped body.

Cell fate analysis in the developing mouse kidney provides an additional insight in support of this view. *Lgr5* expressing cells in the distal S-shaped body have been fate-mapped to multiple distal segments, namely the thick ascending limb of the loop of Henle, the distal convoluted tubule segment, and the macula densa (Barker et al., 2012). The scRNA-seq data shows *LGR5* is expressed broadly in the distal portion of renal vesicles and in S-shaped body nephrons in populations 14, 15, 16, 17, 18 (**Supplementary figure 3D**; strongest in population 16 in the S-shaped body nephron, **Figure 6B**). The human single cell data also predict an overlap with five members of the IRX family of transcriptional regulators, encoded on chromosomes 5 and 16 (Kim et al., 2012). Though, *Irx1* and *Irx3* are necessary for normal nephron patterning in the Xenopus pronephros (Reggiani et al., 2007; Alarcón et al., 2008), it remains unclear whether they serve a role in the formation of the macula densa. In fact, the precise evolutionary origin of the macula densa is not known and it is absent in fish where the Corpuscle of Stannius may instead serve a similar function (Wingert and Davidson, 2008). However, in the human S-shaped body nephron several IRX genes show overlapping expression, with *Irx1* and *Irx2* in a similar configuration in the macula densa in the adult mammalian kidney (**Supplementary figure 9**). Additional mouse cell lineage studies, focused on genes like *PAPPA2*/*Pappa2* restricted to population 18 in the S-shape body and the macula densa of the adult kidney will provide important new insights to the regulation of nephron patterning and morphogenesis, likely shared between mouse and humans.

Cell identities and their behavior are underpinned by complex gene interaction networks. Such networks can be identified by data-driven approaches that integrate thousands of independently generated analyses (Greene et al., 2015). These networks have previously been used to extract genes with nominally significant p-values from GWAS studies. We utilized the power of predicting gene networks and applied this to the resolved cell transcriptomes to generate functional gene networks that are specific for all genes separately within each of the 18 cell-states detected in the early human nephron program. To demonstrate the utility of this approach, we analyzed the gene networks for over 200 genes involved in CAKUT and extracted putative CAKUT genes with strong connections to disease genes. These were further compared against 4 GWAS studies totaling ~1M individuals with data on altered GFR. The predicted CAKUT genes were expressed as predicted and combinations of new and known genes were identified within networks. The integration of existing public data with single cell omic analyses represents a step forward for predicting gene interactions and genes likely responsible for disease.

In summary, our analysis, integrating space and time across across scRNA-seq and threedimensional protein imaging datasets of the human nephrogenic program, generates the most comprehensive view to date of mammalian nephrogenesis. Spatial modelling and transcriptional analysis support an early commitment of some cell types, while others maintain a more extended progenitor capability for distinct regions of the kidney. The data will serve as a valuable resource for mining regulatory pathways generating nephron cell types, facilitating approaches to generate functional nephron structures in vitro. Further, the resulting database (https://sckidney.flatironinstitute.org/) will facilitate identifying potential developmental relationships in gene associations linked to kidney disease.

## Acknowledgments

We thank current and past members of the McMahon lab for helpful discussions and feedback. We would also like to thank Melissa L. Wilson (Department of Preventive Medicine, University of Southern California) and Family Planning Associates for coordinating fetal tissue collection. BH and CA wish to thank Richard Baldock of the MRC Human Genetics Unit, IGMM, University of Edinburgh for supporting this project and providing financial support enabling travel for CA to visit USC and NOL at the outset of the project. We thank Friedhelm Hildebrandt for discussion and informational sources on known CAKUT genes and Pietro Cippa for helpful discussions on GWAS studies.

## Author Contributions

NOL, RS, XC, CA, OGT, and APM wrote the manuscript.

NOL, RS, XC, CA, BH, AS, OGT, and APM designed the experiments.

NOL, RS, XC, RP, AR, GDSB, BH, JC, TT, AW, AWa, AWo, BG, MT, JA, and SR performed experiments, analyses, computational tools, and web-resources.

## Supplementary figure titles and legends

**Supplementary figure 1 – Anatomies for human and mouse nephrogenesis**

(A, B) 3D surface rendering of human and mouse nephrons (blue) at different stages from pretubular aggregate to late S-shaped body stages and the adjacent ureteric epithelium (bone-color), respectively. (C) Immunofluorescently labelled mouse nephrons rendered in 3D. (D) Graph displaying nuclei counts for renal vesicles and pretubular aggregates for human and mouse nephrons. (E) Graph displaying the number of CDH1^+^ and JAG1^+^ cells for renal vesicles for mouse and human. (F) Variance of CDH1^+^ and JAG1^+^ domains for the human and mouse renal vesicles. Horizontal bars in E-F indicate average. Significance (t-test) shown on horizontal lines between categories. (G) 3D rendered human nephrogenic niche (left) and ortho-slices of S-shaped body nephrons showing chirality. (H-I) Composite images of registered nephrons with different protein combinations featuring distal and proximal features. Immunofluorescent labels, features, and developmental stages as indicated on fields.

**Supplementary figure 2 - scRNA-seq analyses of human nephrogenesis**

(A) Work-flow for scRNA-seq experiment from tissue through to data metrics and tSNE plot for whole week 14 data set. (B) tSNE plot displaying replicate week 14 kidneys used in analysis. (C) Violin plots showing mitochondrial gene content, UMI counts, and gene counts for whole week 14 data set. (D) tSNE plot for whole week 14 data set with dotted lines indicating demarcated populations as indicated on plot. (E, F) Dotplots for differentially expressed genes across all week 14 data set clusters and for cell-cycle stage-specific genes. (G) tSNE plot for differentiating nephrogenic cells not in the cell-cycle. (H) Dotplot for same cells as in G using marker genes to identify precursor populations. (I) Dotplot of nephron lineage and cell-type specific marker genes across the 18 cell clusters. (J) SWNE plot for nephrogenic cells not in the cell-cycle (clusters 1-18), data equivalent to that in Figure 3A. PTA: pretubular aggregate. RV: renal vesicle. CSB: comma-shaped body nephron. SSB: S-shaped body nephron. CNT: connecting tubule. LOH/MD: loop of Henle/Macula densa precursor.

**Supplementary figure 3 - scRNA-seq analyses of precursor populations**

(A, B) SWNE featureplots for all nephrogenic cells not in the cell-cycle (clusters 1-18), data equivalent to that in Supplementary figure 2J displaying gene expression for genes activated as described on fields. (C) Dotplot for top differentially expressed genes per cell-types. (D) Dotplot for top 50 differentially expressed genes for each of the 6 populations labelled as S-shaped body cells. Line colors match cluster colors of each precursor. PTA: pretubular aggregate. RV: renal vesicle. CSB: comma-shaped body nephron. SSB: S-shaped body nephron. CNT: connecting tubule. LOH/MD: loop of Henle/Macula densa precursor.

**Supplementary figure 4 - Fine tuning transcriptome origins**

(A-D) Graph-based dimensionality reduction plots of all cell transcriptomes (clusters 1-18). Large circles with numbers and connecting arrows color match cell clusters 1-18 as in Figure 3A. Blue color on plot indicate cells from each cluster. Arrows indicate proposed direction of differentiation. (E) Pearson correlation analysis depicted as dotplot. Numbers on axes correspond to clusters 1-18. Colors relate to Pearson correlation value −1 to +1. NPC: nephron progenitor cell.

**Supplementary figure 5 – Validation of computational analyses of transcriptome relationships**

(A-D) Fluorescent RNA-scope analyses for genes displaying distinct dynamic expression patterns matching those in E-J. Graph-based dimensionality reduction plots in A-D show expression for genes as well as double positive cells in the stated combinations. Genes as indicated on plots. PTA: pretubular aggregate. CSB: comma-shaped body nephron. SSB: S-shaped body nephron. (E-J) Predicted expression profiles for genes generated from the Human Nephrogenesis Atlas. See Figure 3C-D for full details of annotations on lineage-tree plots. Expression levels are indicated in low to high as shown.

**Supplementary figure 6 - Computational analyses of protein patterns**

(A, B) 3D rendering of computed combinatorial protein patterns across different k-values for renal vesicles and S-shaped body nephrons, respectively. k values between 2 and 20 explored. Patterns shown are those which did not belong to background and edge effects. (C) The contribution of each protein to each pattern across k-values as a readout for pattern stability for S-shaped body nephrons. (D) Number of voxels covered by non-background patterns for S-shaped body nephrons across k-values showing erosion of patterns with increased k-values. (E) Registered nephrons rendered in 3D. Immunofluorescent labels as indicated on fields. (F) Graph-based dimensionality reduction plots of all cell transcriptomes (clusters 1-18) showing cells spatially mapped to S-shaped body patterns. Colors correspond to protein patterns as shown in Figure 4D.

**Supplementary figure 7 - Spatially mapping transcriptomes to 3D models**

(A) Dotplot for top 50 differentially expressed genes for the cells mapping to each of the 8 protein patterns. X-axis represents clusters 1-18 from Figure 3A. Colored lines match to protein patterns. (B-D) In situ hybridization of human fetal kidneys showing expression of genes from renal vesicles through to late S-shaped body. RV: renal vesicle. CSB: comma-shaped body nephron. SSB: S-shaped body nephron. SSB+: late S-shaped body nephron. Dotted line in B-D indicates proximal distal axis of nephron. Gene and scales as indicated on fields. tSNE plot of gene expression, and schematic of expression in S-shaped body nephron.

**Supplementary figure 8 – Building functional gene networks for single-cell transcriptomes**

(A) Functional gene network for JAG1 in the connecting tubule of the S-shaped body nephron (cluster 17). (B-C) Two subsets of the hierarchical clustering of 199 CAKUT genes as indicated on Figure 5C. Genes shown are expressed in proximal precursors of S-shaped body and early nephrogenic stages, respectively. Genes plotted against cluster identities 1-18. PTA: pretubular aggregate. RV: renal vesicle. CSB: comma-shaped body nephron. SSB: S-shaped body nephron. CNT: connecting tubule. LOH/MD: loop of Henle/Macula densa precursor. High/Low as indicated. (D) 20 CAKUT genes from Park et al., 2018 as expressed in Park et al. 2018 data from adult mouse kidney and as expressed against cluster identities 1-18 from human data. (E) 199 CAKUT genes - as shown in Figure 5C - contrasted with their expression in the adult mouse kidney data from Park et al., 2018. (F-H) In situ hybridization for *EFNB2* and *TNFRSF11B* on human fetal kidney. Magnified region in Figure 5H is taken from within boxed region in G. Boxed region in F relates to Figure 5E. Scales and stains as indicated. Expression levels and edge weights as indicated by scales on panels. MD: macula densa. RC: renal corpuscle.

**Supplementary figure 9 – Computational precursor-product predictions between human and mouse single cell RNA sequencing data**

(A) Dotplot showing expression of enriched transcription factors in the developing human S-shaped body cell populations (top) and their mouse orthologs in the adult mouse nephron (bottom). (B) Dotplot showing expression of enriched transcription factors in the adult mouse nephron cell populations (top) and their human orthologs in the developing human S-shaped body cell populations (bottom). Cluster numbering on the human S-shaped body corresponds to those in Figure 3D, while those for the mouse nephron match cluster numbers in Ransick et al., 2019. The colors for the clusters correspond to the coloring of Figure 3D and proposed relationships as shown also in model – Figure 6F. Colored arrowheads below the x-axis indicate genes with conserved expression domains between human and mouse nephrons according to proposed relationships.

**Supplementary figure 10 – Exploring precursor-product relationships in the S-shaped body**

(A) Immunofluorescently labelled human S-shaped body nephron. Dotted area indicates HNF4A^+^ domain. White arrowhead indicates autofluorescence from endothelial cells. Schematic depicts proposed relationship between proximal precursors and proximal convoluted tubule (PCT). (B) tSNE plot for week 17 zone1 (cortex) and zone 2 (medulla) scRNA-seq data. (C) Dotplot for cell marker genes using week 17 data. (D) Feature plots showing expression of indicated genes and tSNE plot indicating the zone1 and zone2 cell origins. Black arrowhead indicates *NOS1^+^/PAPPA2^+^* population. (E) Dotplot for genes highly correlating with *PAPPA2* in the week 17 data. Green marking indicates cluster and genes highly enriched in the *PAPPA2^+^* cluster. (F) Feature plots showing expression of indicated genes.

**Supplementary figure 11 – Macula densa precursor expression signatures**

(A-C) Immunofluorescently labelled human S-shaped body nephrons – white arrowheads indicate macula densa precursors. Dotted line marks the proximal distal axis. Stains as indicated on fields.

## Tables with titles and legends

**Supplementary table 1 Differential gene expression in whole week 14 kidneys**

Differential gene expression analyses between all cells in replicate 1 and replicate 2 of the week 14 human fetal cortex cell isolations as shown in Supplementary figure 2A.

**Supplementary table 2 Differential gene expression during nephrogenesis**

Differential gene expression analyses between cells in clusters 1-18 as shown in Figure 3A.

**Supplementary table 3 Differential gene expression between mapped transcriptomes**

Differential gene expression analyses between cells mapping to patterns 1-8 as shown in Figure 4D.

**Supplementary table 4 CAKUT genes and their neighbor genes**

Sheet1. 285 CAKUT genes from literature. 272 of them are included in the network. Sheet 2. 199 CAKUT genes are active in at least one cell type (expressed in >10% cells). Normalized gene expression value per gene per cell type is shown. Sheet 3. Summary for connectivity of 272 CAKUT genes in each of the 18 cell type-specific functional networks and number of differential genes identified per cell type using p-value 0.05 for cutoff. Sheet 4. Differentially expressed CAKUT neighbor genes’ and validation analyses against GWAS.

## STAR Methods

### Contact for Reagent and Resource Sharing

Further information and requests for resources and reagents should be directed to and will be fulfilled by Nils O. Lindström, Olga Troyanskaya and Andrew P. McMahon.

### Experimental Model and Subject Details

#### Animal studies

All animal work performed in this study was reviewed and approved by the Institutional Animal Care and Use Committees (IACUC) at the University of Southern California. All work adhered to institutional guidelines. Embryos were recovered at embryonic day 15.5 following timed matings of Swiss Webster mice. The sex of embryos was not known. For *Nos1* lineage-tracing, crosses were set as previously described (Riquier-Brison et al., 2018). In brief, *Nos1Cre^ERT2^* (Nos1tm1.1(cre/ERT2)Zjh; Taniguchi et al., 2011) crossed with a conditional florescent protein reporter *Rosa26^mTm^* (B6.129(Cg)-Gt(ROSA)26Sortm4(ACTB-tdTomato,-EGFP)Luo/J; Muzumdar et al., 2007). Postnatal day 5 pups were injected with tamoxifen and harvested either 1 day or 20 days later.

#### Human samples

Consented, anonymized, human fetal kidney tissue was obtained from elective terminations following review of the study by Keck School of Medicine of the University of Southern California’s Institutional Review Board. Kidney samples ranging in age from 14 to 17 weeks of gestation were provided by collaborators at Family Planning Associates. Gestational age was determined per guidelines specified by the American College of Obstetricians and Gynecologists using ultrasound, heel to toe, and crown to rump measurements following published Carnegie Stages (O’Rahilly and Müller, 2010; O’Rahilly et al., 1987; Strachan et al., 1997). The sex of the specimen was not reported. Consented samples were received immediately after elective terminations and transported from the Children’s Hospital of Los Angeles on ice at 4°C in 10% fetal bovine serum, 25mM Hepes, high glucose DMEM (SigmaAldrich).

### Immunofluorescent labelling of mouse and human kidneys

Whole embryonic day 15.5 Swiss Webster kidneys were dissected, decapsulated in 1xPBS by standard dissection tools and fixed for 20 min at 4°C in 4% PFA without shaking. Human kidneys were carefully decapsulated in 1xPBS, placed on surgical gauze in 10cm Petri dish submerged in 1xPBS. Blunt-ended forceps were used to hold the kidney and 3 mm thick cortical slices were sliced off manually using a scalpel. Human kidney slices were fixed 4% PFA in 1xPBS for 45 minutes at 4°C without shaking. Human and mouse kidneys were subsequently washed with several rounds of 1xPBS.

For preparation for immunofluorescent staining slices and kidneys were blocked for 1hr in 1xPBS with 2% SEA Block and 0.1% TritonX100 at 4°C with gentle movement. The primary antibodies were resuspended in blocking solution and kidney tissue incubated in primary antibody at 4°C with gentle movement for 48 hr. Samples were washed up to 8 hr through several rounds of 1xPBS with 0.1% TritonX100. Secondary antibodies were resuspended in blocking solution and tissue incubated at 4°C with gentle movement for 48 hr. Washing steps for primary antibody were repeated. Tissue was counterstained in 1 *μ*g/ml Hoechst 33342 in 1xPBS for 2 hr prior to final wash in 1xPBS. For dehydration steps prior to clearing steps, the tissue was passed through a series of increasing concentrations of methanol (50%, 75%, 100%) 1 hr for each step. To clear tissue the dehydrated tissue was immersed in 50% Benzyl alcohol, Benzyl benzoate (BABB) / 50% methanol solution for a 1 hr. Full clearing was achieved in 100% BABB. The tissue was stored at 4°C in the dark.

| Antibody | Vendor | Cat no. |
| --- | --- | --- |
| anti-Ap-2-α | Cell Signaling | 3215S |
| anti-Cdh1 | BD Transduction laboratories | 610182 |
| anti-Cdh6 | R&D | AF2715 |
| anti-Cldn5 | Novus Biologicals | NB120-15107 |
| anti-Emx2 | R&D | AF6470 |
| anti-ErbB4 | R&D | MAB1131 |
| anti-Foxc2 | R&D | AF6989 |
| anti-Hnf1b | Santa Cruz | sc-22840 |
| anti-Hnf4a | R&D | MAB4605 |
| anti-Irx1 | SIGMA | HPA043160 |
| anti-Jagged1 | R&D | AF599 |
| anti-Krt8 | DSHB | TROMA-1 |
| anti-Lef1 | Cell Signaling | 2230 |
| anti-Lhx1 | Santa Cruz | sc-19341 |
| anti-Mafb | R&D | MAB3810 |
| anti-Mafb | Santa Cruz | sc-10022 |
| anti-Mecom | R&D | MAB75061 |
| anti-Pappa2 | R&D | AF1668 |
| anti-Pax2 | Biolegend | 901001 |
| anti-Pax8 | Abcam | ab189249 |
| anti-Podxl | R&D | AF1658 |
| anti-Pou3f3 | Thermo Scientific | PA5-64311 |
| anti-Pou3f3 | Novus Biologicals | NBP1-49872 |
| anti-Sall1 | R&D | PP-K9814-00 |
| anti-Six2 | Mybiosource | MBS610128 |
| anti-Six2 | ProteinTech | 11562-1-AP |
| anti-Sox9 | Abcam | ab185230 |
| anti-Tfcp2l1 | Novus Biologicals | AF5726 |
| anti-Tfap2a | Cell Signaling | 3215 |
| anti-Tfap2b | Cell Signaling | 2509S |
| anti-Wt1 | Abcam | ab89901 |

### *In situ* hybridization and RNA-scope

RNAscope probes were designed and purchased (Advanced Cell Diagnostics) and reactions performed on human kidney cryosections according to manufacturer’s recommendations. The Multiplex Fluorescent Reagent kit v2 was used. Probes were used for *SLC39A8, TMEM72, COL4A4*, *LAMP5*, *HNF4A*, *EFNB2*.

All *in situ* hybridization stains were performed on human kidney cryosections as described in detail at the GUDMAP website: https://www.gudmap.org/Research/Protocols/McMahon.html and as used previously on human sections (Lindström et al., 2018b).

### Image acquisition

Imaging of cortical slices (3D) was performed on a Leica SP8 using a 40X objective (40x/1.30 Oil HC PL APO CS2). 1024×1024 images were captured at 0.35 μm optical intervals with a 1.5x zoom for human samples and 2x zoom for mouse kidneys. Channels were captured sequentially. Images were captured at 12-bit depth over a 0-4095 range. Care was taken to ensure voxels were not saturated.

### Image Processing Pipeline

A mainly automated image processing pipeline was developed to process, register and combine protein localization assay images from both human and mouse renal objects allowing quantitative comparison of protein abundance. The pipeline, written in Python, makes extensive use of both the ANTs and Woolz image processing systems. ANTs was used for elastic and some affine image registration; Woolz for all other image processing tasks including the affine registration of S-shaped body-stage nephrons.

The image processing pipeline, ANTs and Woolz are freely available from the GitHub repositories:

Registration pipeline: https://github.com/ma-tech/RenalObjectRegistrationPipeline

ANTs: https://github.com/ANTsX/ANTs

Woolz: https://github.com/ma-tech/Woolz

### Image Data Preparation for 3D Mapping

Confocal images were organized into assays, with each assay composed of the set of segmented images obtained from a single nephron. In all cases each assay contained the DAPI and JAG1 channel images for reference, but the assays also contained a selection of two other channels from the set CDH1, LEF1, SALL1, SIX2, SOX9, TROMA1, WT1, LHX2, PAX2, HNF1B, FOXC2, MECOM, EMX2, TFAP2A, POU3F3, ERBB4, PAPPA2, CDH6, MAFB, and CLDN5. Four classes were used to group the assays of nephron objects, namely human and mouse renal vesicles and human and mouse s-shaped body stage nephrons.

The confocal images were rescaled using rescaling factors of 0.4103 within the image plane and a factor of 0.9889 across planes to achieve cubic voxels. Composite DAPI and JAG1 channel rescaled images were created by combining the voxel values of these two channel with a JAG1:DAPI weighting ratio of 4:1. The composite images were created to combine the broad coverage of the DAPI channel with the directional coverage of the JAG1 channel. A Gaussian blurred version of each of the rescaled images was also generated for visualization and registration using an isotropic Gaussian filter with a sigma value of two voxels.

### Image Registration

For each of the nephron object classes a representative assay was selected as the basis of the class model and each assay was then registered to the corresponding model using a multistage registration process. First a general affine transform was established to align each assay object with its model using the Gaussian blurred combined DAPI - JAG1 images. While ANTs was used for renal vesicle affine registration, ANTs proved unreliable when establishing an initial affine registration of S-shaped bodies due to their morphology, so landmarks manually placed using WoolzWarp were used to identify the proximal, medial, and distal compartments. From these landmarks, least squares affine transforms were computed using Woolz. With the transformations that oriented and scaled assay images to their corresponding models established, ANTs was used to elastically register combined (but not blurred) DAPI – JAG1 images to their models using a cross-correlation similarity metric. The resulting affine and elastic transforms were then used to register all channels of all assays to their models. With all assays of all models aligned, model images were subsequently composed from the combined registered single channel assay images.

### Extracting 3D protein patterns from registered models

For each protein with imaging data, we first found the average protein expression at each voxel across all images for the protein in both the renal vesicle and S-shaped body image sets. We then normalized across all voxels so that the intensities of each protein had a mean 0 and standard deviation of 1 within both the renal vesicle and the S-shaped body datasets. Selecting the set of 18 or 19 proteins with imaging data for renal vesicle and S-shaped body, respectively, we performed k-means clustering on each set of voxels, treating each voxel as an 18 or 19 - dimensional point in genomic space. We use the k-means implementation in the stats R package for clustering, and tested values of k ranging from 2 to 20. We then visualized the resulting spatial patterns using Fiji, manually removing patterns that corresponded to background noise.

### Capture, sequencing, and processing of single-cell RNA transcriptomes

Single cells were isolated from two replicate week 14 kidneys as previously described (Lindström et al., 2018a). The nephrogenic niche was dissociated by enzymatic digestion by placing the decapsulated but intact whole kidneys in collagenase A/ pancreatin, an enzymatic cocktail described elsewhere (Brown et al., 2015). The kidneys were kept incubated at 37°C in a nutator and shaken at 450rpm to release cells from the nephrogenic niche over a 40-50min period. Released cells were stained for DRAQ5+ (ThermoFisher Scientific) and DAPI (ThermoFisher Scientific) to identify live and intact and dead cells, respectively. DRAQ5+/DAPI-cells were selected by FACS. Per repeat, 7000 cells were input into the 10X Chromium system and processed for single-cell library construction as per 10x Genomics instructions and as we describe previously (Lindström et al., 2018a). Cell transcriptomes were sequenced by Hi-Seq (Illumina) in 8 separate runs with ~ 120,000 reads per cell. 24,254 cells were sequenced to a mean of 2644 genes per cell (**Supplementary figure 2A**).

Quality control, mapping (to GRCh37.p13) and count table assembly of the library was performed using the CellRanger pipeline version 2.1 (as consistent with 10x Genomics guidelines) and as described in our previous work (Lindström et al., 2018a).

### Computational isolation of nephrogenic lineage

To begin analyzing the cell transcriptomes we assessed and filtered transcriptome qualities using Seurat (Satija et al., 2015). Cells with more than 3000 genes per cell, fewer than 5% of reads mapping to mitochondrial genes, and a Good-Turing estimate greater than 0.7 (Good, 1953) were kept for downstream analyses. We deliberately set the gene content to 3,000 in order to keep only cells that displayed very rich information. 8,316 remaining cells were clustered using the Seurat R package. We ran Principal Component Analysis on the dataset and used 39 principal components based on the JackStraw test (p < 0.05) and clustered the cells using the Seurat FindClusters function with 39 PCs and default remaining parameters. We found 39 resulting clusters (**Supplementary figure 2D**). Based on the differential expression test (FindAllMarkers function, bimod test) and expression of marker genes we determined that 24 of the clusters, a total of 6,667 cells, were nephrogenic (**Supplementary figure 2E**).

Genes linked to cell-cycling (e.g. *TOP2A* and *MKI67*) were expressed strongly in several clusters (**Supplementary Figure 2E**) and on closer scrutiny against genes with known periodicity during cell-cycle phases (Dominguez et al., 2016), clusters containing cells in the cell-cycle were categorized based on cycle-stage (**Supplementary figure 2F**). Non-cycling nephrogenic cells (2893 cells) were selected for downstream analyses (nephrogenic lineage clusters in **Supplementary figure 2F**). To maximally resolve precursor populations, the nephron progenitor cells were initially removed (*CITED1^+^, SIX2^+^*), and differentiating cells (clusters 20,27,19,25,18,17,26,5) reiteratively reclustered and identified based on marker gene expression and gene enrichment analyses (**Supplementary figure 2G, H**). The subclustering was repeated as described above (using 30 principal components, and clustering explored using a range of nearest neighbour values ranging between 4 and 30), analyzing the differentially expressed genes between resulting clusters for segregation of known marker genes of the predicted subdomains. We identified 3 clusters displaying a very early nephron signature, which we termed pretubular aggregate clusters. Similarly, we identified 5 clusters that consisted of transcriptomes from renal vesicle cells, and 6 clusters from S-shape body nephrons (see **Results** section for details of annotations). Note that cluster numbers in **Supplementary figure 2G, H** do not correspond to those in **Supplementary figure 2C-F**). Resolved transcriptomes of differentiating cells were then remerged with the nephron progenitors (**Figure 3A**) and cell relationships explored by Similarity Weighted Nonnegative Embedding (SWNE) (**Supplementary figure 2J**) (Wu et al., 2018) which showed three main trajectories corresponding to distal, proximal, and podocyte fates, with the clusters distributed along the trajectories in an order reflecting developmental progression. The SWNE projection was consistent with the tSNE plot (**Supplementary figure 2J** vs. **Figure 3A)**. Based on marker gene analysis (**Supplementary table 2**) and work outlined in the **Results** section, the cluster annotations 1 to 18 were assigned to the resolved clusters based on developmental relationships.

### Single cell RNA sequencing data availability

The human week 14 scRNA-seq data presented in this work is GEO accession number GSE139280.

Zonal human week 17 scRNA-seq data was obtained as described (Tran et al., 2019) GSE127344.

Adult mouse nephron scRNA-seq data was obtained as described (Ransick et al., 2019) GSE129798.

### Computing cell relationships from single cell RNA sequencing data

Cell relationships were computed by graph-based dimensionality reduction (Menon et al., 2018) and Zhou and Troyanskaya https://www.biorxiv.org/content/10.1101/2020.04.12.022806v1).

### Mapping single cells to voxel data

To map single cells to protein patterns, we imputed genes for the proteins with confocal imaging data using the SAVER algorithm (Huang et al., 2018). For each gene, we normalized its expression across the single-cell data. Genes were scaled to give each gene mean expression of 0 and standard deviation 1 across all cells. We next selected the fourteen genes with matching confocal imaging data that were highly expressed in the S-shaped body scRNA-seq clusters (*EMX2, MECOM, POU3F3, SOX9, PAX2, PAPPA2, HNF1B, LHX1, ERBB4, KRT8, CDH6, JAG1, MAFB*, and *WT1*). We further selected the 8 protein patterns identified (see **Results**) and added a null pattern with zero expression of all proteins. We computed the Euclidean distance between the vector of highly expressed genes and the corresponding vector of pattern centers for each cell and each pattern center. Each cell was assigned to the closest pattern center. We identify genes differentially expressed in the cells assigned to each pattern using the FindAllMarkers function in the Seurat package.

### Annotating human gene functional relationships

To infer functional relationship between genes, first we extract cell type-naïve gene functional relationships from the Gene Ontology database. Based on a set of 618 expert-selected GO biological process terms (GO evidence codes: EXP, IDA, IPI, IMP, IGI and IEP), pairs of genes that were co-annotated to the same terms after propagation were treated as positive (i.e., functionally related) examples. Gene pairs not co-annotated to any of these terms were considered as negative examples. This resulted in a cell type-naive gold standard of 472,239 functionally related gene pairs (positive examples) and 11,430,296 potentially unrelated pairs (negative examples).

### Predicting cell type-specific gene functional interactions

To learn cell type-specific gene functional interactions, we used Seurat to identify cell typespecific genes, which are actively (fold change > 0) and significantly (Wilcoxon test, adjusted *p*-value < 0.05) expressed in each of 18 cell types. For each cell type, we labelled each gene pair as cell type-specific if both genes were differentially expressed in the current cell type (C,C), or non-specific if at least one gene is not differentially expressed in current cell type (but differentially expressed in other cell types) (C,C’) or (C’,C’). Overlapping cell type-specific gene pair labels with the above positive (+) or negative (-) functional relationship gold standard we obtained six classes of gene edges as follows:

· (C+C): if both genes are differential genes in the current cell type and functionally related in the gold standard;
· (C+C’): if one gene is differential in the current cell type and the other gene is a marker gene from any other cell type, and the two genes are functionally related;
· (C’+C’): if both genes are from any other cell type and functionally related;
· (C-C): if both genes are differential genes in the current cell type and not functionally related;
· (C-C’): if one gene is differential in the current cell type and the other gene is a marker gene from any other cell type, and the two genes are not functionally related;
· (C’-C’): if both genes are from any other cell type and not functionally related.

To predict cell type-specific functional relationships between genes, C+C gene pairs representing functional relationships of cell type-specific genes were labeled as positive examples while equal proportions of edges in the other five classes were labeled as negative examples. For each cell type, we trained a naive Bayesian classifier by integrating labeled cell type-specific genes pairs with 1,541 genome-scale data sets (including 1533 human gene expression data sets from GEO, interaction data from BioGRID, IntAct, MINT and MIPs, transcription factor co-regulation from JASPAR, genetic perturbation (c2:CGP) and microRNA target (c3:MIR) from MSigDB) (Wong et al., 2018). Using the learned classifier, for each gene pair, we predicted its cell-type specific functional relatedness in the current cell type.

### Identifying genes functionally associated with CAKUT disease

To identify genes functionally associated with CAKUT disease, we integrated fetal kidney scRNA-seq data with cell type-specific gene functional networks and glomerular filtration rate (GFR) GWAS studies. Specifically, for each cell type, we selected genes actively expressed in this cell type based on our scRNA-seq data (in more than 10% cells). Then, given the built cell type-specific gene functional network, we selected genes whose interactions to 285 CAKUT disease genes were significantly stronger than interactions to 4,029 non-CAKUT disease genes in the OMIM database (Wilcoxon test, *p*-value <0.05). Using four GFR GWAS traits, we filtered the above gene list using significant GWAS SNPs (*p*-value < 0.05).

### Additional resources

The Human Nephrogenesis Atlas (NephMap) displays scRNA-seq data in three displays (1) as a lineage tree with expression levels averaged across the cluster, (2) as an anatomical diagram of the nephrogenic niche using the same values as in the lineage tree, and (3) as a tSNE plot. The atlas enables individual or multi-gene searches for gene networks within each cell-type as defined in the 18 clusters. Further, 3D rendering views are presented for the predicted protein patterns as well as the protein abundance for each of the proteins used for the construction of the S-shaped body nephron.

