## Supplementary figures and images for "Spatial Transcriptional Mapping of the Human Nephrogenic Program"

### Supplementary figure 1

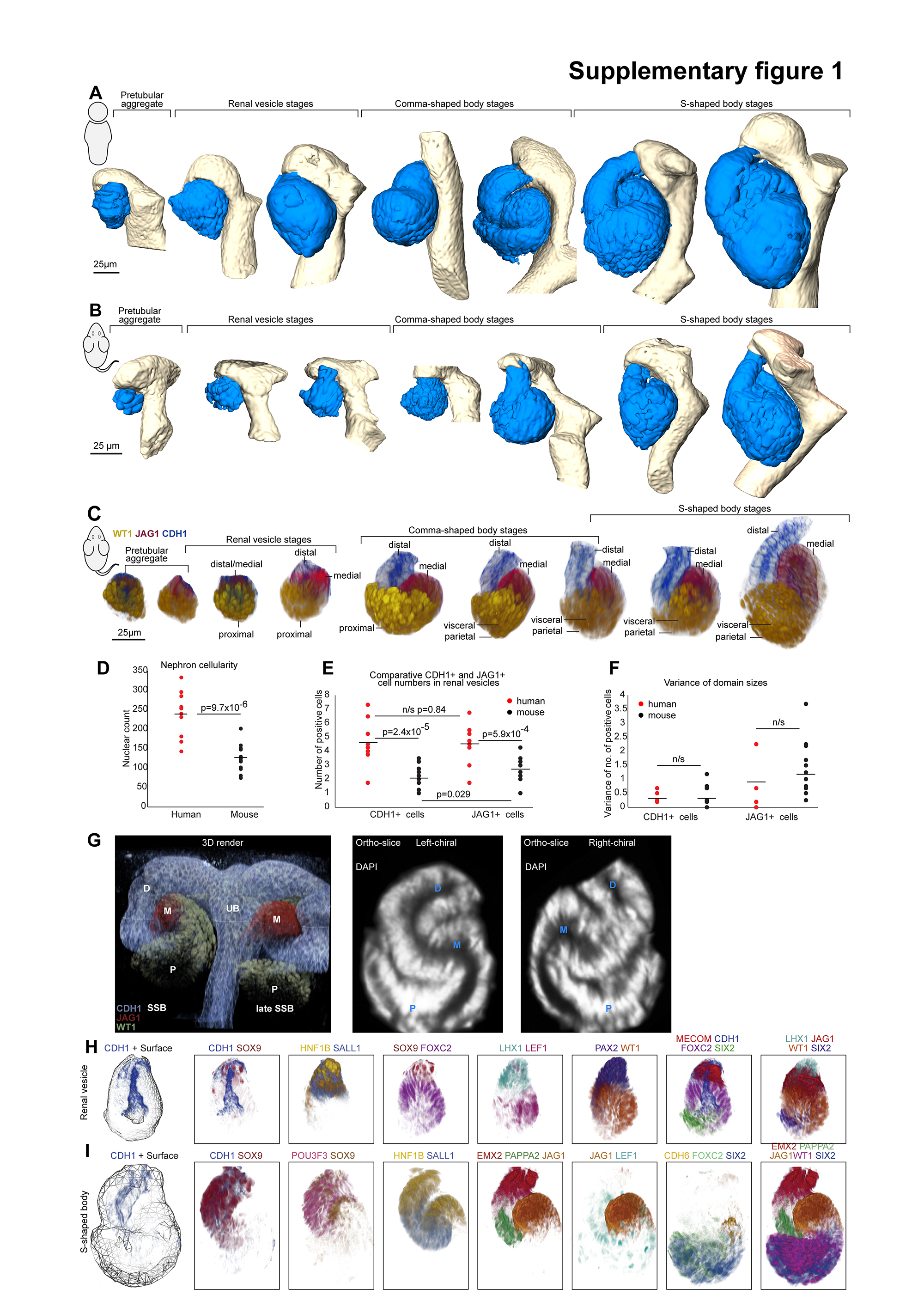

### Supplementary figure 2

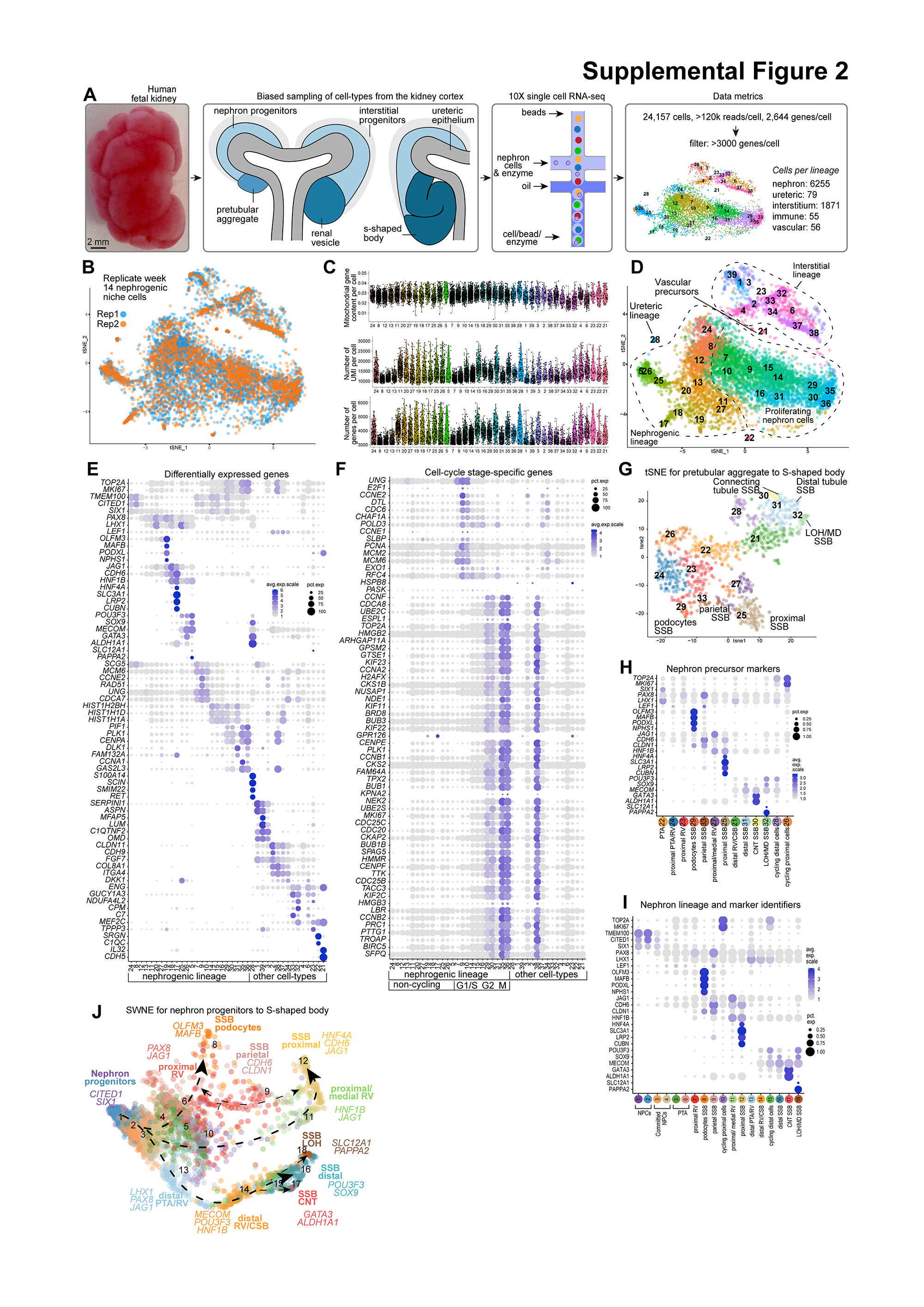

### Supplementary figure 3

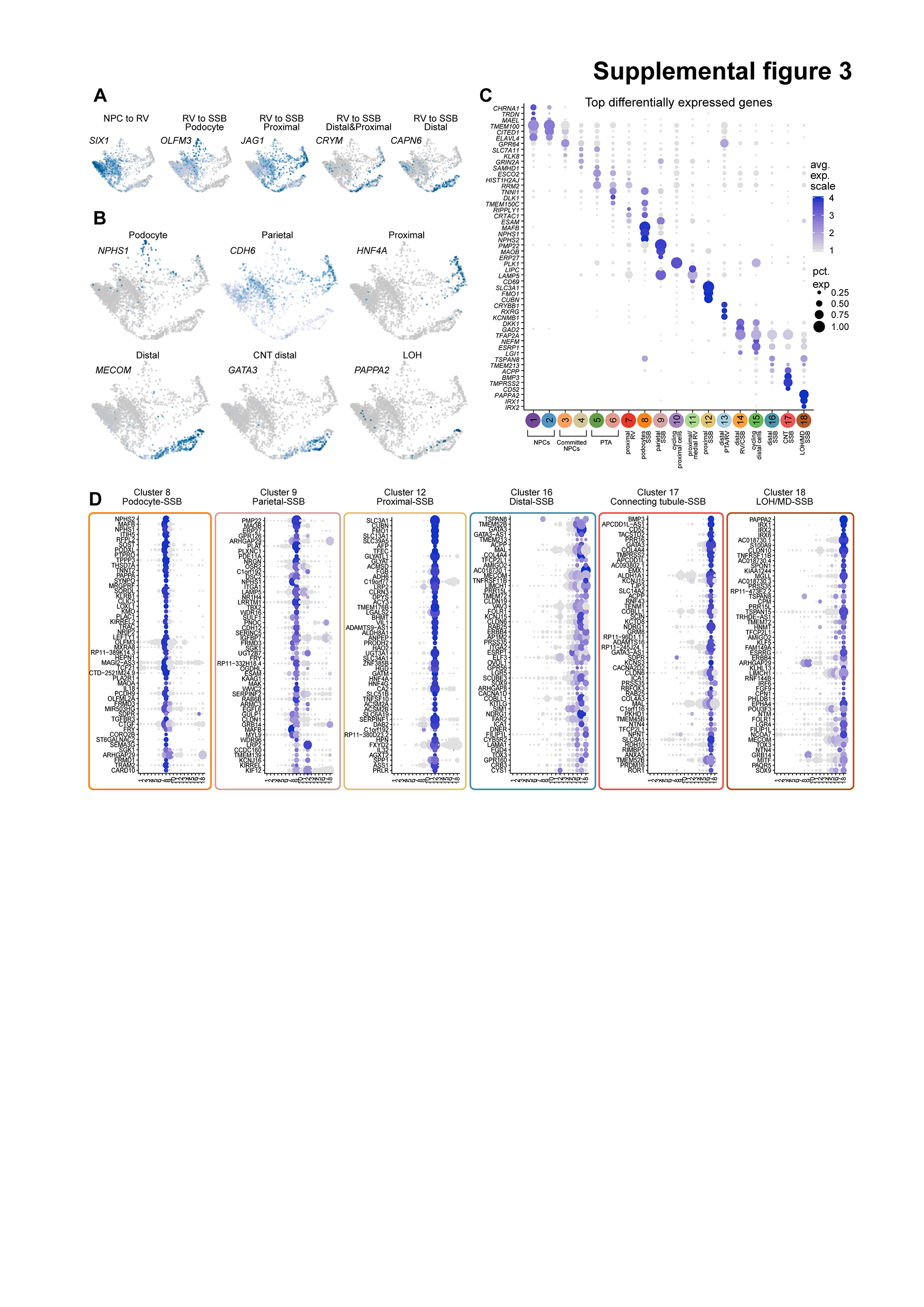

### Supplementary figure 4

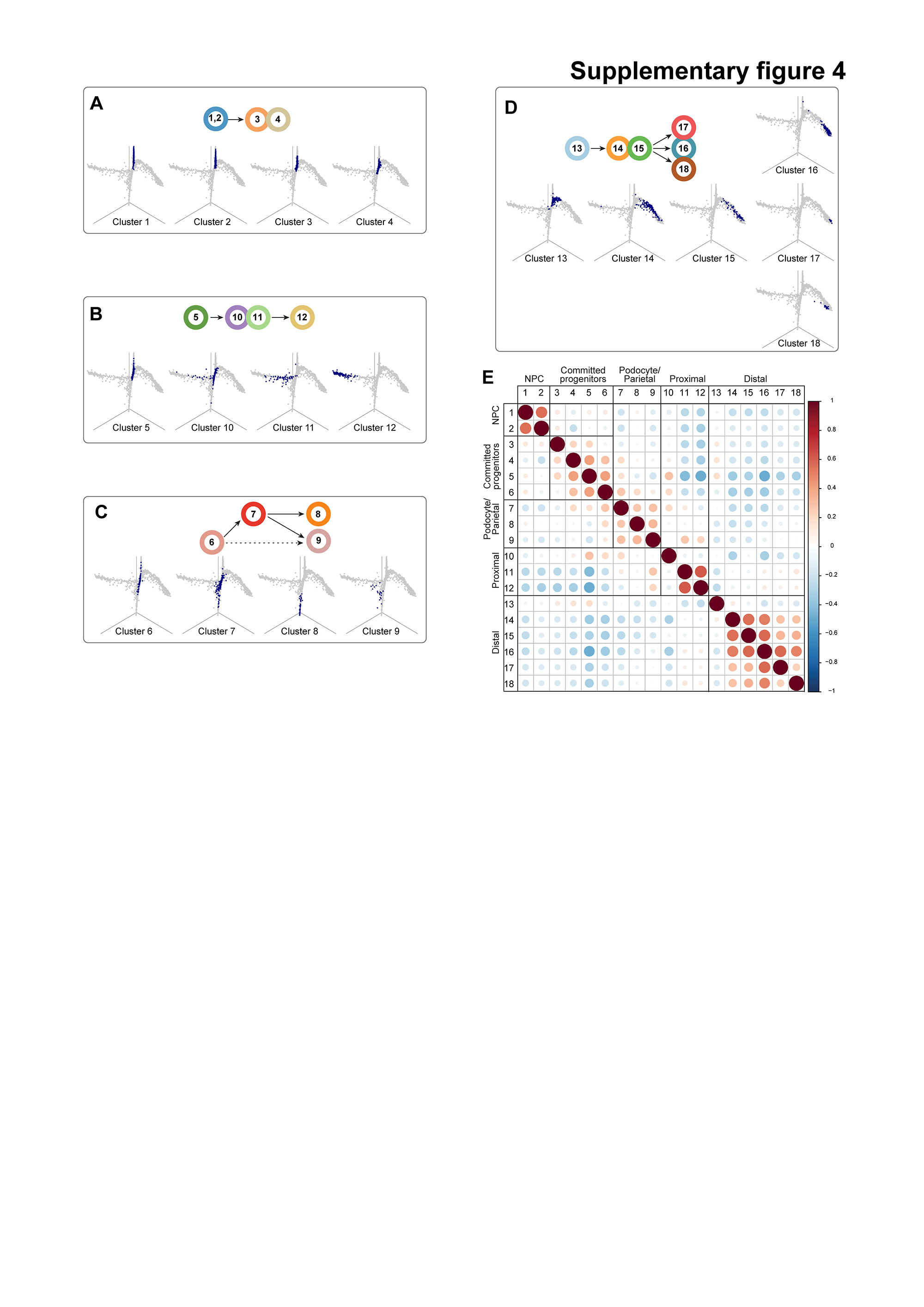

### Supplementary figure 5

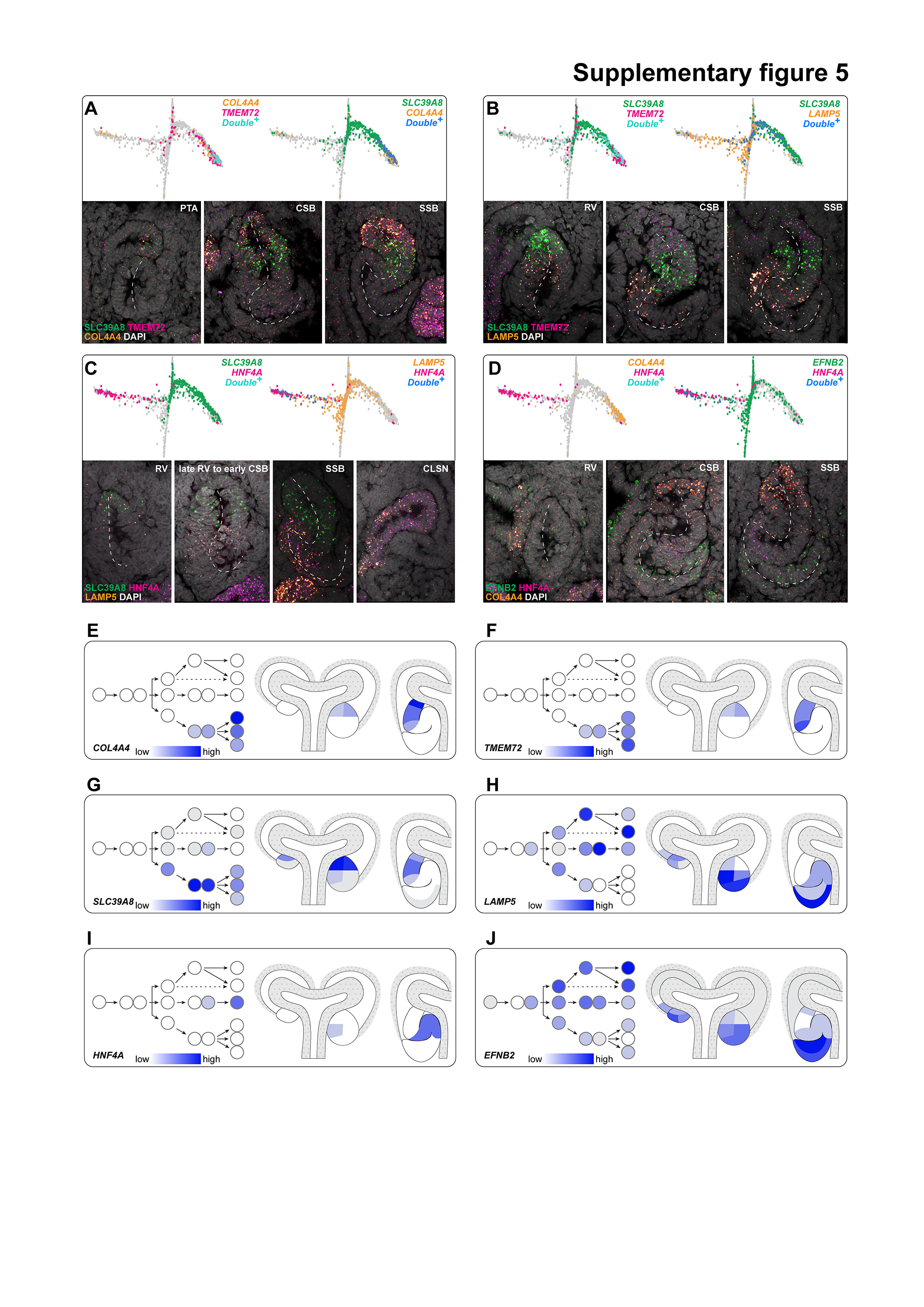

### Supplementary figure 6

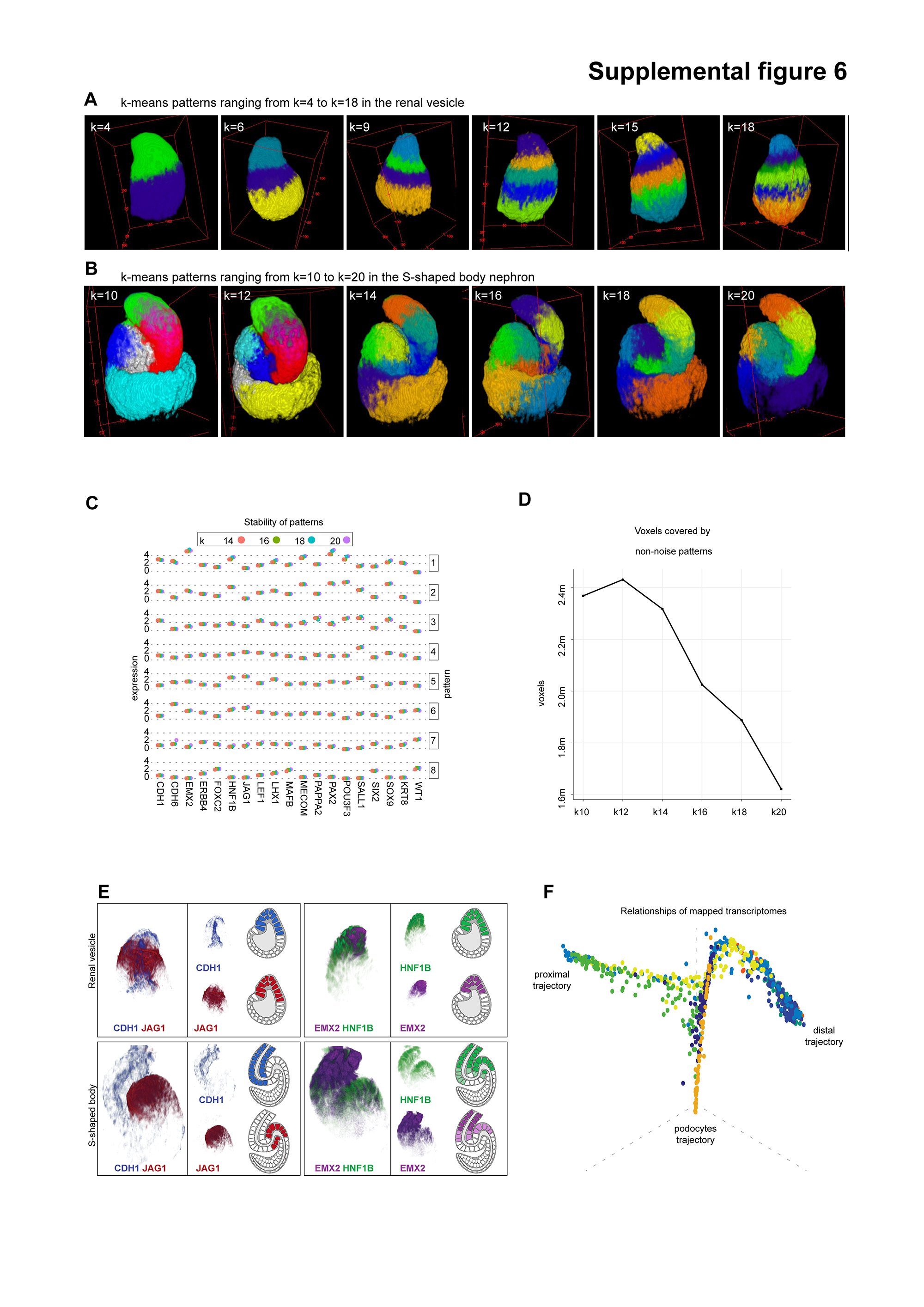

### Supplementary figure 7

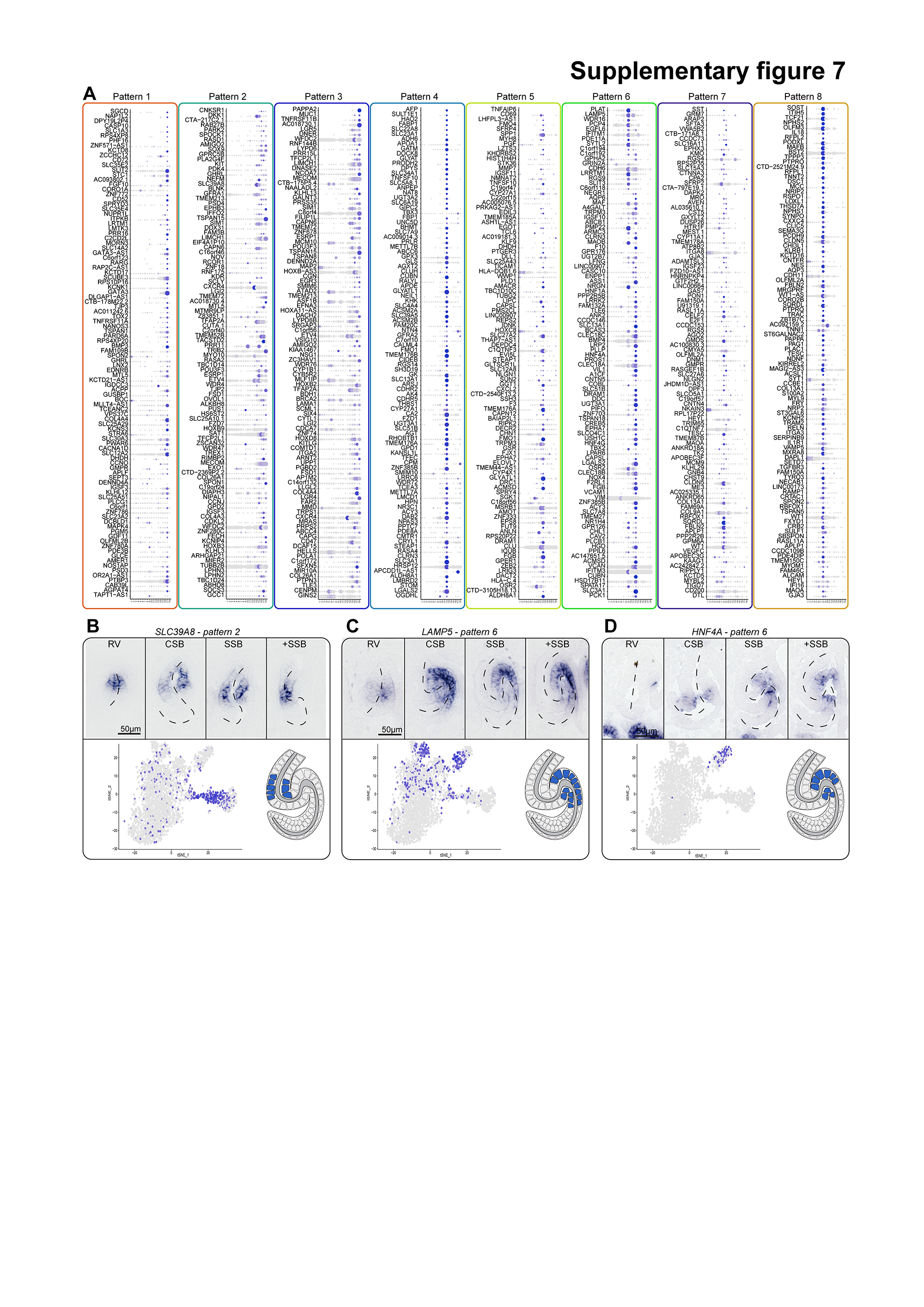

### Supplementary figure 8

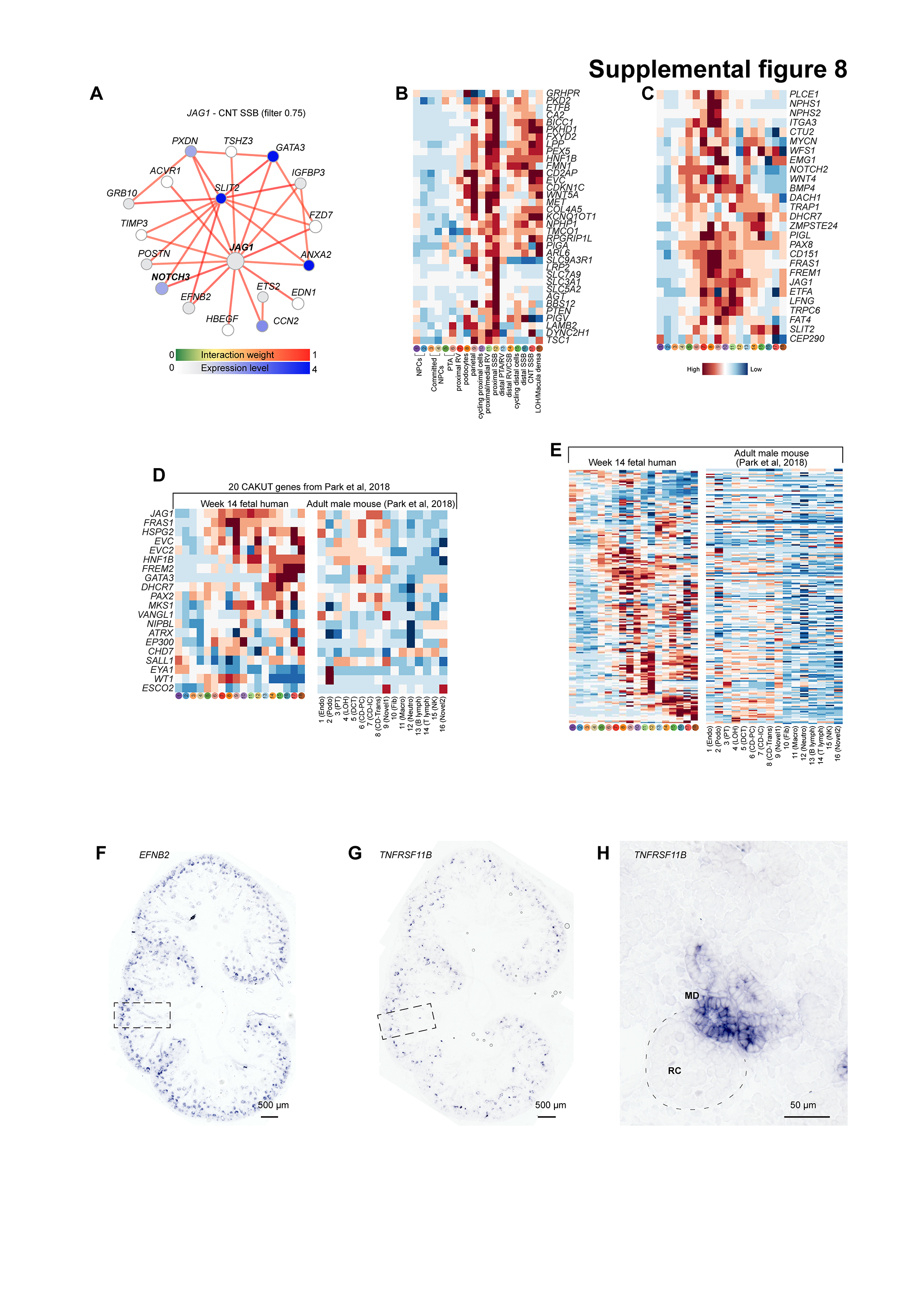

### Supplementary figure 9

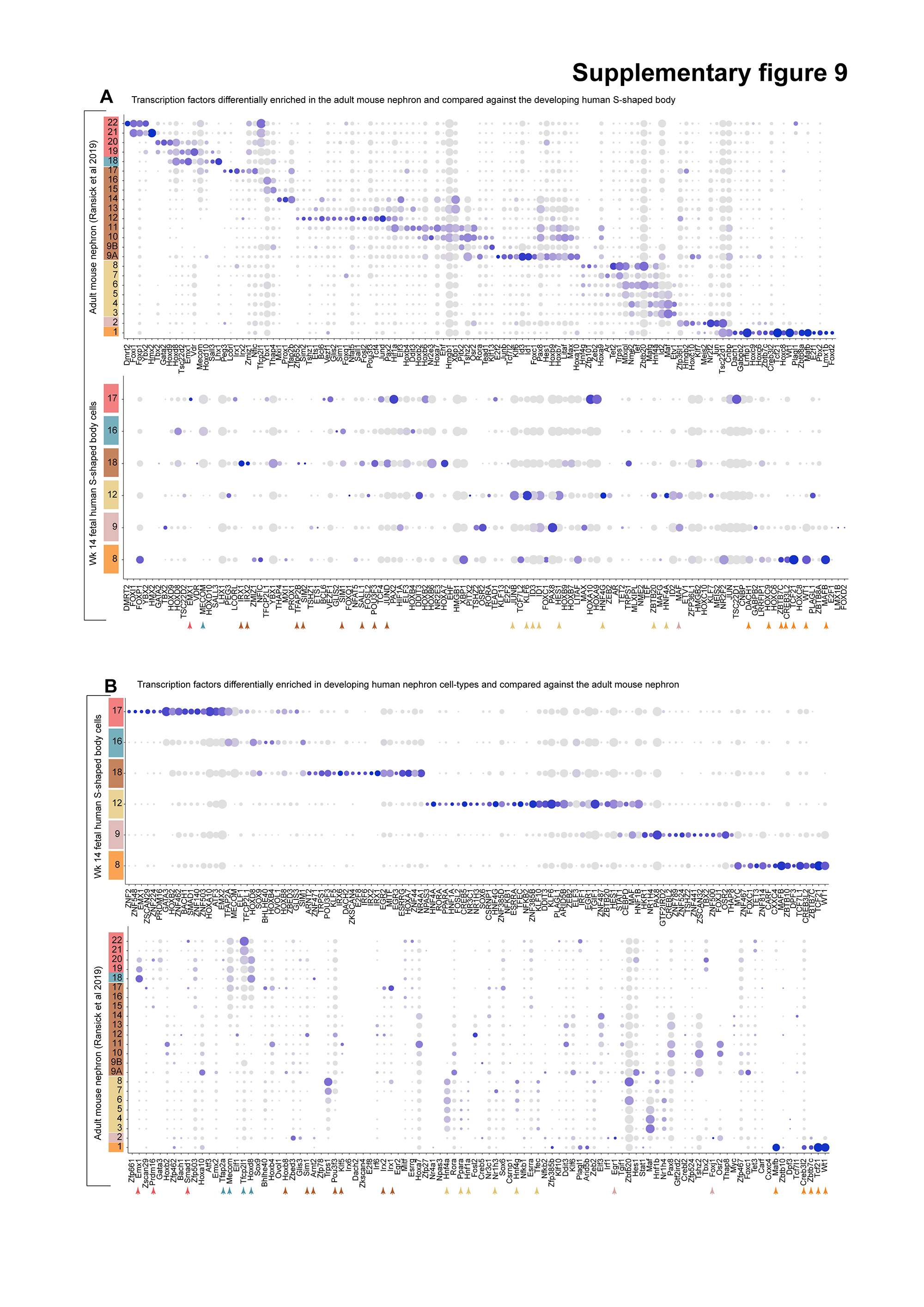

### Supplementary figure 10

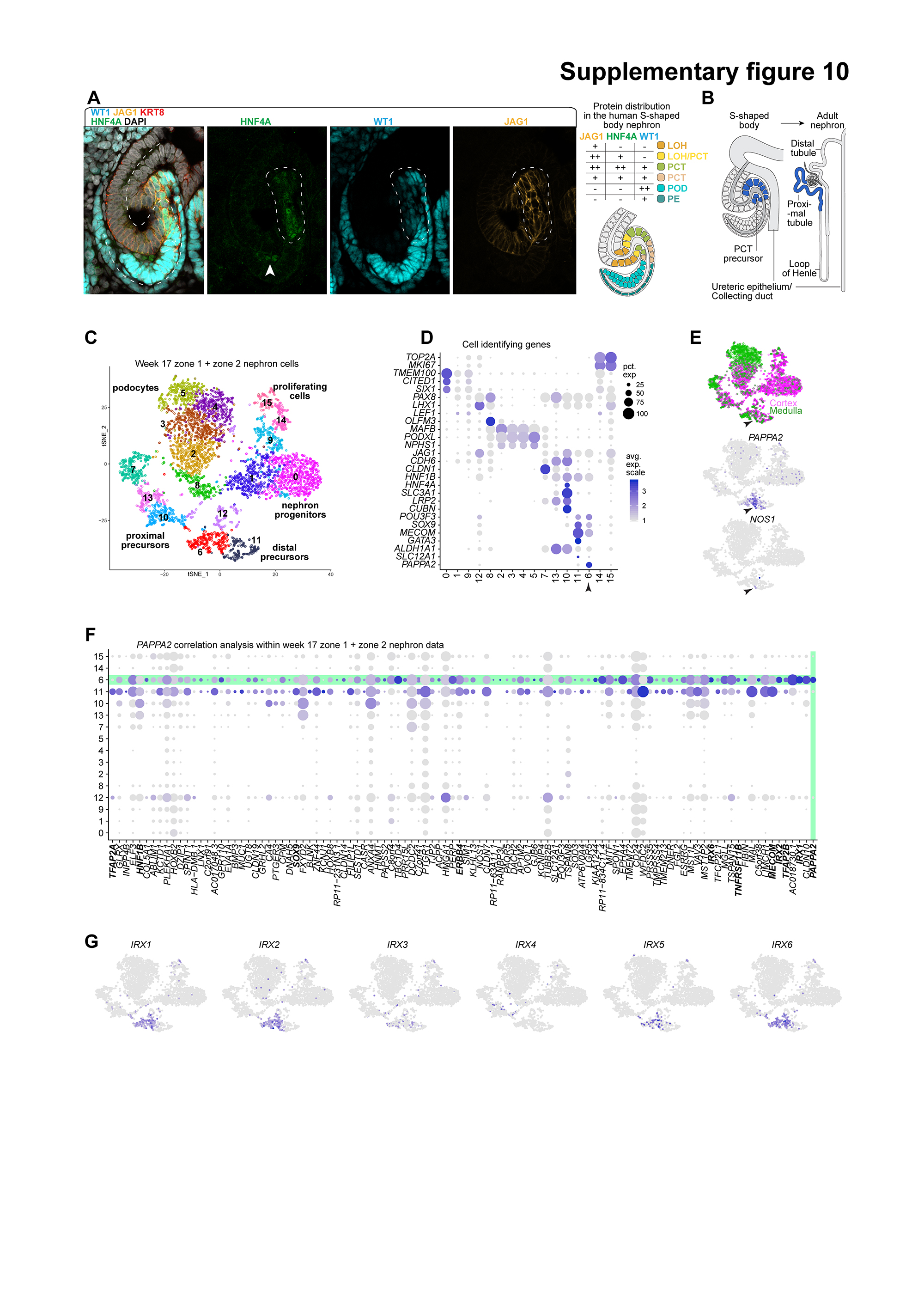

### Supplementary figure 11

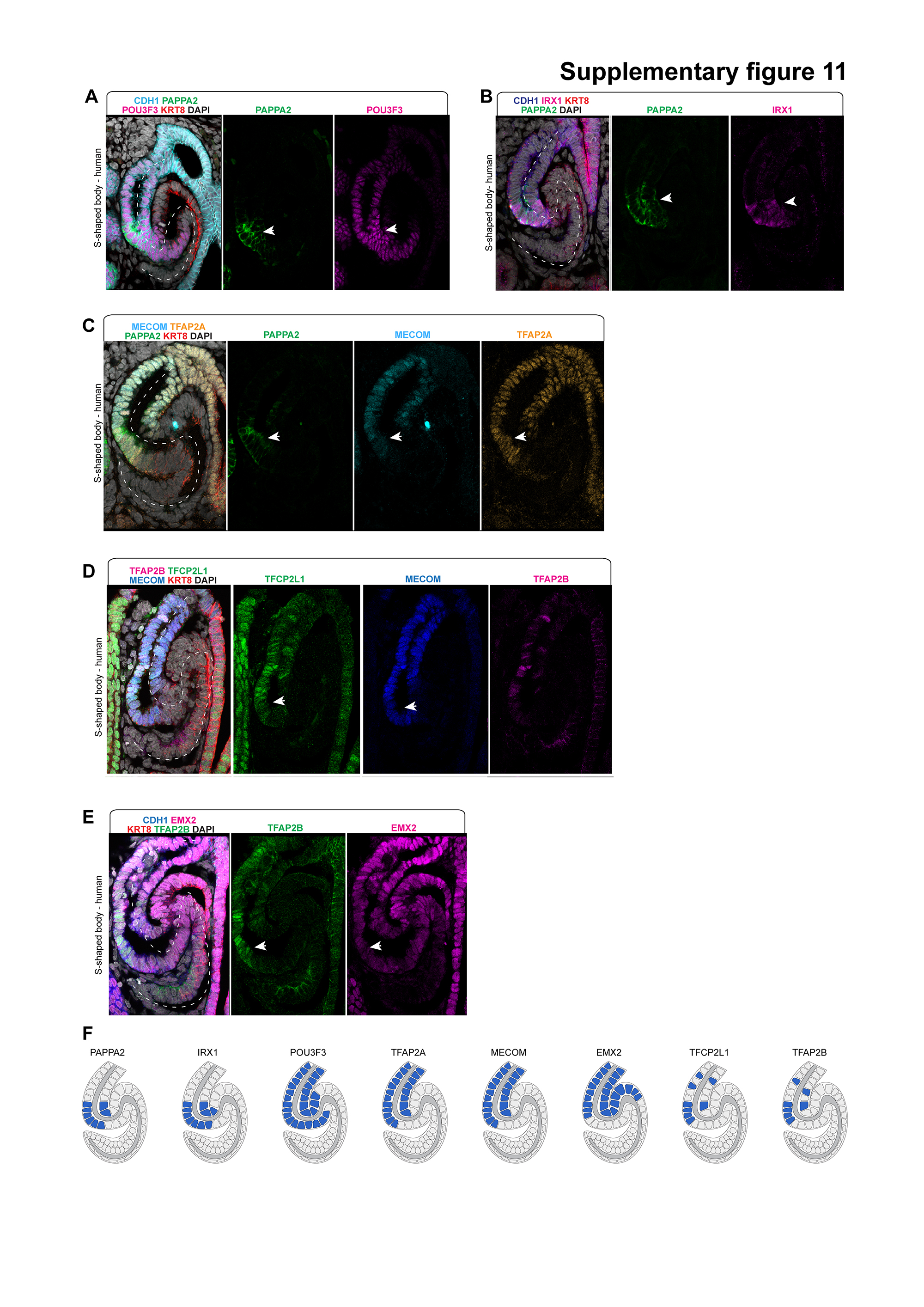
